# Neural mechanisms for resilient sensory computation under metabolic stress

**DOI:** 10.64898/2026.10.29.761790

**Authors:** Sebastian H. Castedo, Zahid Padamsey, Nathalie L. Rochefort, Rémi Monasson

## Abstract

Food-restricted mice can solve visual discrimination tasks despite reduced synaptic currents and increased noise resulting from cellular changes in the primary visual cortex. Combining neural recordings and modelling, we study the population structure of the noise and show it is mainly concentrated along task-irrelevant directions in high-dimensional population activity, thereby preserving coding on the low-dimensional stimulus manifold. In addition, we predict that food restriction shortens decision times, conserving energy but limiting the evidence accumulation needed for fine discrimination.

## Metabolic constraints increase noise correlations

Calorie restriction induces energy-saving cellular changes in the primary visual cortex (V1) of mice (Fig. 1a)^1^. In this food-restricted state, ATP consumption is reduced as a result of decreased synaptic currents. Concomitantly, neurons maintain their average firing rate by increasing input resistance and baseline membrane potential. This compensation increases subthreshold variability in the membrane potential, mostly due to slow synaptic input noise (~85%), and, to a lesser extent, fast cell-intrinsic noise (~15%)^1^. Remarkably, while single neurons are less informative about stimuli, the average decoding performance of the whole neural population is largely maintained, which points to collective error-correcting properties of the system (Extended Data Fig. 1c). Accordingly, food-restricted animals accomplish visual discrimination tasks with the same behavioural performance as control animals (fed ad libitum), unless the tasks are particularly challenging (Fig. 1a)^1^.

**Fig. 1.**
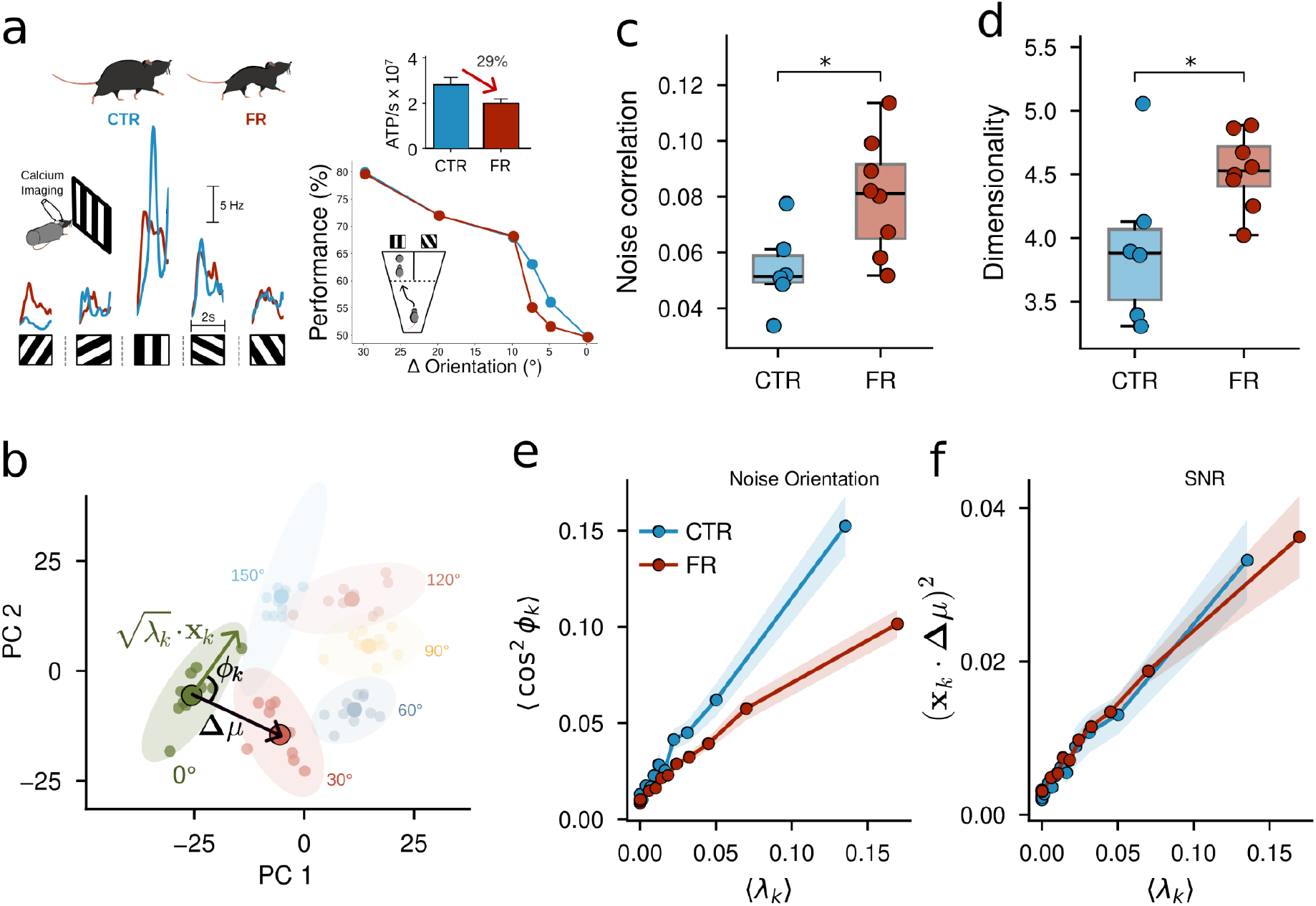
Food restriction reshapes population noise while preserving stimulus discriminability. **a:** A summary of results from ^1^. Mice were food restricted to lose 15% of their body mass. Examples of visual responses of two deconvolved layer 2/3 neurons imaged with two-photon calcium imaging in awake mice during the presentation of drifting gratings. A loss in orientation selectivity and an increased trial-to-trial variability were found in food-restricted (FR) animals compared to control (CTR) animals (bottom left), accompanied by a strong reduction in ATP consumption rate (top right). In a Y-shaped water-maze task requiring mice to distinguish grating orientations, food-restricted mice perform similarly to controls for coarse discriminations, but worse for fine discriminations (bottom right). **b:** Principal component analysis of the stimulus-averaged population activity from a particular control animal reveals a two-dimensional circular manifold of responses. For each stimulus, dots show mean activity vectors for each trial and ellipses show the trial-to-trial noise covariance projected into the same global activity PC plane. For a highlighted stimulus orientation, the leading noise mode is shown as the vector 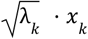. The signal vector, Δµ, connects the mean population responses of two stimuli, while ϕ denotes the angle between the signal and noise vectors. Equation (1) relates each mode’s contribution to discrimination to these geometric terms. **c:** Noise correlations are higher in food-restricted (FR) animals than controls (CTR). Pearson correlations were calculated between the trial-to-trial residual activity during the presentation of a grey screen (see Methods). Points represent individual animals; boxes show the median and interquartile range. Exact two-sided animal-label permutation test, *p* = 0. 023. Correlation time courses are found in Extended Data Fig. 1a. **d:** Food restriction increased the effective dimensionality of trial-to-trial population variability (see Methods). Points represent individual animals; boxes show the median and interquartile range. Two-sided permutation test on log-transformed animal-level participation ratios, *p* = 0. 035. **e:** Squared cosine of the angle between the signal and noise mode, ϕ, against the noise covariance eigenvalue, λ_*k*_, with both quantities averaged across animals and stimuli. In food-restricted animals, the noise increases relative to controls, but the largest noise modes are less aligned along the signal vector. **f:** The squared overlap between the signal vector and the noise mode is plotted against the eigenvalue with both quantities averaged across stimuli. By Eq. (1), their ratio gives the mode’s SNR contribution; the similar slopes indicate that SNR is preserved.

The impact of single-neuron noise on population coding depends on how fluctuations are shared across neurons and whether they obscure differences between stimuli in the high-dimensional space of the activity (Fig. 1b). The effect of variability on discrimination depends on its alignment with the signal vector, the difference between the mean population responses to two stimuli (gratings of different orientations). Noise along the signal direction obscures this difference and impairs discrimination. Noise orthogonal to it does not^2^. To investigate this population structure, we analysed two-photon calcium imaging of V1 neuronal populations of control and food-restricted mice. We calculated pairwise correlations between neurons from trial-to-trial fluctuations in neuronal responses to repeated presentations of the same stimulus ^2^ (see Methods). These *noise* correlations increased under food restriction (Fig. 1c). This increase disappeared once the food-restricted mice had recovered their body mass after two weeks of ad libitum access to food (Extended Data Fig. 1a).

## Geometric reorientation of noise covariance modes compensates for noise increase

How can an increase in noise correlations, which is generally detrimental for decoding in V1^3,4^ (Extended Data Fig. 1c), have little impact on the behavioural performance of food-restricted mice? To answer this question, we decomposed the high-dimensional noise covariance matrix into its eigenvectors *x*_*k*_, and eigenvalues λ_*k*_, which quantify the variances of the noise along the modes *x*_*k*_.

Eigenvalues increased on average, with the largest proportional increase found in weak modes (Extended Data Fig. 1b). The additional variability was therefore distributed across more population directions, flattening the covariance spectrum and increasing the effective dimensionality of the population variability (Fig. 1d). The simultaneous increase in correlations and dimensionality is unexpected, as stronger correlations typically confine population activity to fewer dimensions.

We then calculated the angles ϕ_*k*_ between the noise and signal directions for both control and food-restricted mice. For both conditions, we observe a linear relationship between cos^2^(ϕ_*k*_) and λ_*k*_ (Fig. 1e), in agreement with a previously reported phenomenological power-law fit^5^. However, the slope is lower for food-restricted animals (red line), indicating a reorientation of dominant noise modes away from the signal direction. After the food-restricted mice had recovered their body weight, we observed a return to the control baseline (in mice fed ad libitum) (Extended Data Fig. 2a). Thus, under food restriction, the increased noise is rendered benign through reorientation along directions that are less important for signal discrimination.

To quantify the effect of reorientation on performance, we computed the signal-to-noise ratio (SNR) for optimal linear discrimination associated with each mode *k* (see Methods),

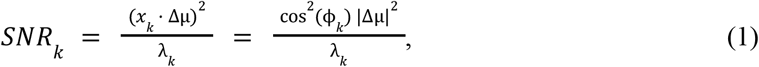

where |Δµ| is the norm of the signal vector. We observed that the SNR is almost perfectly maintained across the different modes between food-restricted and control conditions (Fig. 1f). In other words, the decrease in the slope of Fig. 1e is compensated by a corresponding increase in the squared norm of the signal vector. This result is independent of the type of stimuli, as the same effect is observed when assessing the neuronal selectivity to both the spatial frequency of drifting gratings and the individual frames of a natural movie (Extended Data Fig. 2b,c). Our findings were corroborated by a computational estimation of the full noise covariance structure, using Gaussian processes^6^, which bypasses the limited number of available trials (Extended Data Fig. 3a).

Altogether, these results show that the increased noise associated with food restriction, which reduces orientation selectivity at the single-neuron level (Fig. 1a), is mitigated by population-level mechanisms, including the reorientation of the noise modes.

## Circuit model and linear-response theory

To explain the reorientation of Fig. 1e and the SNR maintenance of Fig. 1f, we developed a model of neuronal populations, analysed it theoretically, and tested it using simulations. Figure 2a shows a diagram of a neural network, with recurrent synaptic connections *W*_*ij*_. Each neuron *i* is subject to an external input *I*_*i*_ (θ), where θ denotes the stimulus (e.g. grating orientation), a slow input noise ξ_*i*_, and a fast cell-intrinsic noise η_*i*_. The membrane potential at time *t* is given as

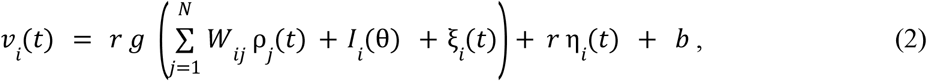

where ρ_*i*_ (*t*) = *F*[*v*_*i*_ (*t*)] is the instantaneous firing rate with a static transfer function *F*[⋅]. Consistent with cellular changes reported in ^1^, the food-restricted state increases the input resistance with respect to controls by the factor *r* = 1.22, reduces the synaptic conductance by the factor *g* = 0.71 and depolarizes the baseline membrane potential by Δ*b* = 6 mV.

**Fig. 2.**
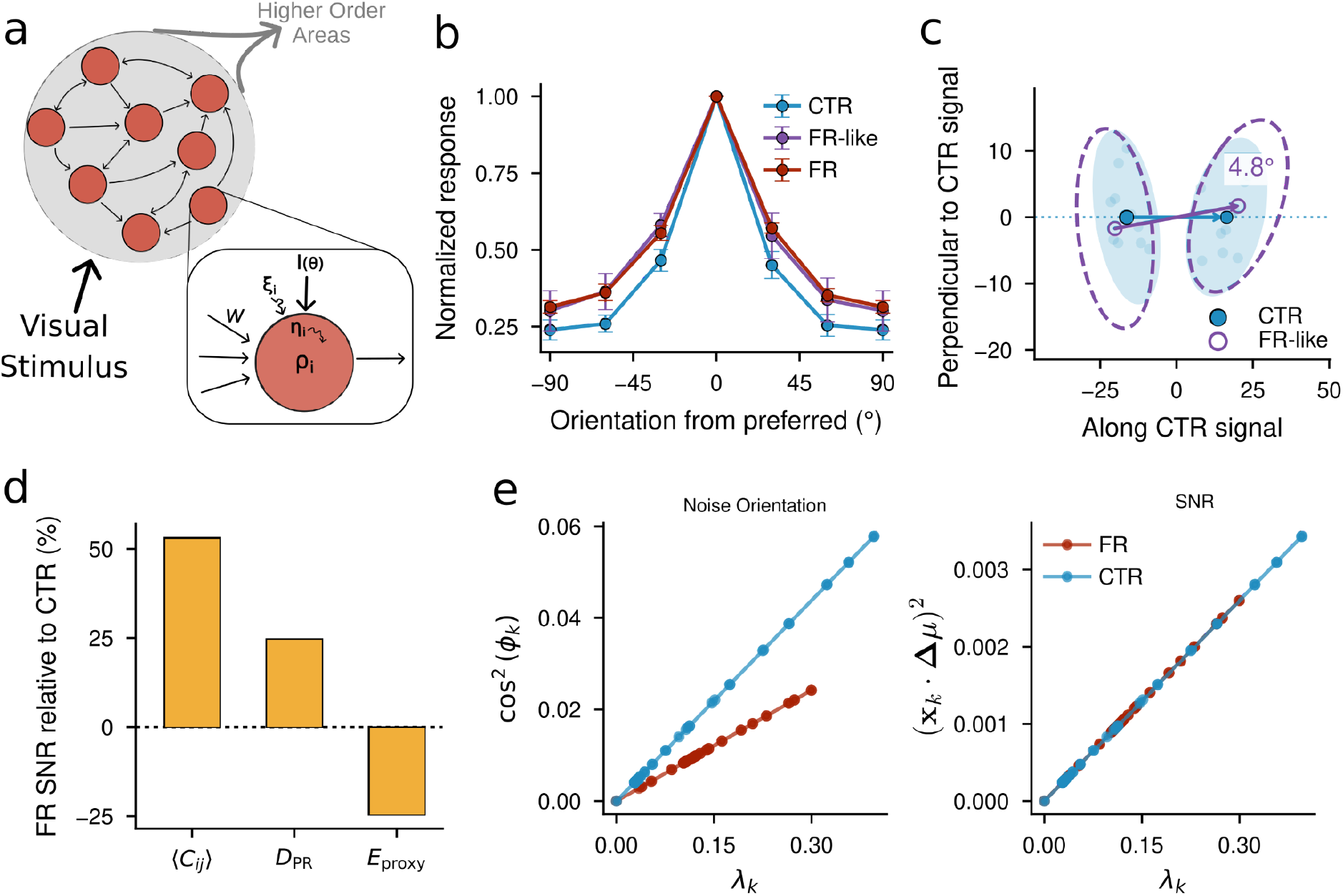
A circuit model links cellular adaptations to preserved population coding under metabolic constraints. **a:** Generic V1 circuit model encoding population responses to stimuli and projecting to higher-order areas. A point neuron is zoomed in on, showing recurrent connections with weights *W*, external input from stimuli, *I*, input noise, ξ, and cell intrinsic noise, η. **b:** We applied a transformation using linear response theory in Eq. (3) (see Methods) to the control data that mimics the food-restricted state. The experimentally found orientation-selective tuning curves are transformed and reproduce the food-restricted data, which indicates that the theory accurately captures metabolic changes to the network. **c:** The experimentally found covariance ellipses are transformed in a similar way. We see that the means are further apart, the ellipses are larger, and they are angled more orthogonally with respect to the control signal vector. We project onto the plane containing both the CTR and FR signal vectors, choosing the horizontal axis along the CTR signal and the vertical axis perpendicular to it. **d:** Mean-field simulations of a circuit model (see Methods), as depicted in Fig. 2a, show how food-restricted cellular changes increase mean correlations and dimensionality, while reducing energy use, consistent with the data in Fig. 1. **e:** Mean-field simulations of a neural network model show that the angle between the signal and the noise directions increases during food restriction and SNR is maintained across the different modes, as seen in the data in Fig. 1.

As the stimulus is slightly varied, so are the external inputs *I*_*i*_ and the neural activities ρ_*j*_. A matrix object, the linear response *R*, describes how the system reacts to perturbations by the external input and can allow us to formulate a theory for the discrimination of different stimuli from each other. A mean-field calculation of *R* as a function of *W* and of the parameters *r, g*, which takes into account the average over the fast noise, is reported in Supplementary Information. In particular, the expression for the change in the signal vector using linear response is given, in vector notation, by

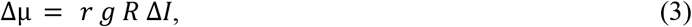

where Δ*I* is the change in the external inputs received by the neurons. Perturbations to the activity induced by the noise, ξ, are mediated by the same filter, *R*, as changes due to external stimuli (see Methods). This is because they are both pre-synaptic inputs and enter the equation at the same level. Setting aside for now the contribution of the fast cell-intrinsic noise, which accounts for only 15% of the total noise in Eq. (2)^1^, the expression for the noise covariance matrix is

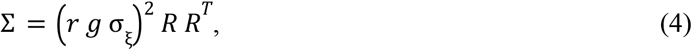

where σ_ξ_ is the standard deviation of the noise.

### Theory captures food-restricted changes to the network

This approach allowed us to explain the changes in the network response, *R*, and thus several features, such as the widening of the tuning curves and the reorientation of the noise. We first obtained *R* using Eq. (4) from the measured noise covariance for control mice and used Eq. (3) to relate the changes in the stimuli (grating angles) to the changes in the recorded neural activity, and then extracted the value of Δ*I*. We then transformed *R* by replacing the cellular parameters for the control mice with those of food-restricted animals, while keeping Δ*I* unchanged (see Methods). Figure 2b shows that the transformed orientation tuning curves broaden, in excellent agreement with the tuning curves observed in V1 neurons of food-restricted mice.

As a further check, we transformed the covariance ellipses and population means of control mice in Fig. 2c, finding good agreement with the food-restricted data (see Methods and Extended Data Fig. 4). The transformation reproduces both the increase in noise, indicated by larger ellipses, and the increase in signal, indicated by greater separation between population means. It also captures the reduced alignment between the signal and the dominant noise directions. This arises because the elevated baseline and larger membrane fluctuations make previously weakly driven neurons more responsive, while neurons that were already strongly responsive change comparatively little. Consequently, fluctuations along previously weak population directions grow more than those along the dominant directions, redistributing variability across the population (as seen in Extended Data Fig. 1b). The same changes also strengthen the stimulus response along these previously weak directions, reducing its alignment with the dominant noise fluctuations. This is further discussed in the Supplementary Information.

## Overall signal-to-noise ratio is unchanged in the food-restricted state

Although changes in the cellular parameters *r* and *g* between control and food-restricted conditions strongly affect both signal and noise covariance, they leave the signal-to-noise ratio *SNR* of each mode *k* unchanged. Using Eq. (3) and Eq. (4), we see that the global signal-to-noise ratio, 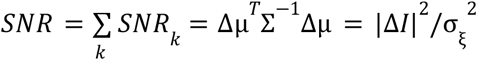, is preserved under food restriction, as found in the *in vivo* data in Fig. 1f. This theory applies generally provided that input amplitudes and fluctuations are unchanged, which was found to be the case ^1^. Crucially, this cancellation does not depend on the particular changes in the network response, *R*, induced by food restriction. Because the stimulus signal and input noise enter through the same pathway, they undergo the same transformation, leaving the SNR unchanged irrespective of how *R* changes.

To provide further evidence that our model fully recapitulates the observations made under food restriction, we numerically simulated the stationary states of the network described in Fig. 2a under time-independent inputs (see Methods). Figure 2d shows that the mean correlation and dimensionality increase, whereas estimated energy expenditure decreases. We used the total presynaptic current as a proxy for energy expenditure because synaptic transmission constitutes the largest energetic burden on neurons^7^. As expected, simulations confirmed that the largest noise modes are more orthogonal to the signal vector during food restriction and that the SNR is maintained (Fig. 2e). We also showed that linear response theory holds for a large range of input differences (Extended Data Fig. 5), thus indicating that our results are valid in most regimes.

The linear-response theory therefore makes two distinct predictions. First, changes in the network response, *R*, account for the broadening of tuning curves, the increase in correlations and dimensionality, and the reorientation of noise relative to the signal. The model also reproduces the linear relationship between signal–noise overlap and noise eigenvalue observed in Fig. 1f. Second, despite these substantial changes in population activity, the stationary SNR remains unchanged, irrespective of how *R* is modified. Food restriction can therefore substantially reshape the geometry of neural activity without reducing the discriminability of the population code.

However, this invariance of the stationary SNR does not imply that changes in the network response have no consequences for discrimination. In particular, the theory does not yet explain the reduction in behavioural performance observed for small differences in grating orientation. Behavioural decisions are made using neural activity accumulated over finite time windows, where rapid neuronal fluctuations become important. We therefore turn to the dynamics of population coding at shorter timescales, where the cancellation underlying the stationary SNR no longer necessarily holds.

### Signal-to-noise ratio in food-restricted state is not maintained on short-time scales

To investigate population coding over finite time windows, we performed *microscopic* simulations of the network in Fig. 2a (see Methods). We modelled V1 as a continuous attractor neural network (CANN) through an appropriate choice of the interaction matrix *W* ^8^. Each neuron stochastically fires or remains silent at each time step, with a probability determined by its membrane potential in Eq. (2). We find that the SNR is maintained under food restriction only when neuronal activity is averaged over sufficiently long time windows (greater than 150 ms; Extended Data Fig. 6a). At shorter timescales, rapid fluctuations in neuronal output are not averaged out, and the covariance is no longer accurately described by Eq. (4).

To update the expression for Σ in the presence of intrinsic stochasticity in the neural activity, we extended the linear response theory to take into account the fast noise η_*i*_ (*t*) in Eq. (2). As described in Supplementary Information, we obtained

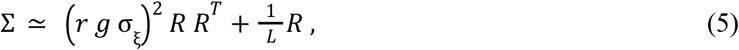

where *L* is the ratio of the duration of the time window over the correlation time of the cellular noise. The additional contribution, linear in *R*, is reminiscent of the fluctuation-dissipation relationship in statistical physics connecting covariance (Σ) and response (*R*). It decreases as 1/*L* because independent fluctuations are averaged together. The presence of multiple contributions to Σ can also be found in recent literature describing information-limiting correlations and bounds on sensory encoding ^9, 10^. Since the signal is quadratic in *R* and the updated covariance is now no longer purely quadratic, the SNR is worse for food restriction, as shown in Extended Data Fig. 6a. We verified the correctness of the new expression in the insets of Extended Data Fig. 6a, where the empirical covariance calculated from the simulation results is in excellent agreement with Eq. (5). For larger averaging windows, the largest eigenvalues in the data follow the quadratic scaling in Eq. (4) as expected, but deviations can be observed for small eigenvalues reconstructed with the Gaussian process technique (Extended Data Fig. 3b).

### Food restriction favours faster decisions at the expense of accuracy

The temporal dynamics of the SNR computed from V1 data over 50 ms windows were plotted in Fig. 3a. Values from food-restricted mice were marginally better for the first 500 ms after stimulus onset, after which SNR values from controls were higher; the dynamics in the food-restricted animals that recovered their body weight closely matched those of controls (Extended Data Fig. 7). To isolate the contribution of noise reorientation, we recalculated the food-restricted SNR using signal-noise angles from control animals, ϕ_*k*_^*CTR*^, while retaining the food-restricted signal magnitudes and noise eigenvalues (see Eq. (1)). Without reorientation, the transient SNR advantage over controls disappeared (Fig. 3a). This indicates that the reorientation identified in Fig. 1e is essential for the early enhancement of discriminability under food restriction.

**Fig. 3.**
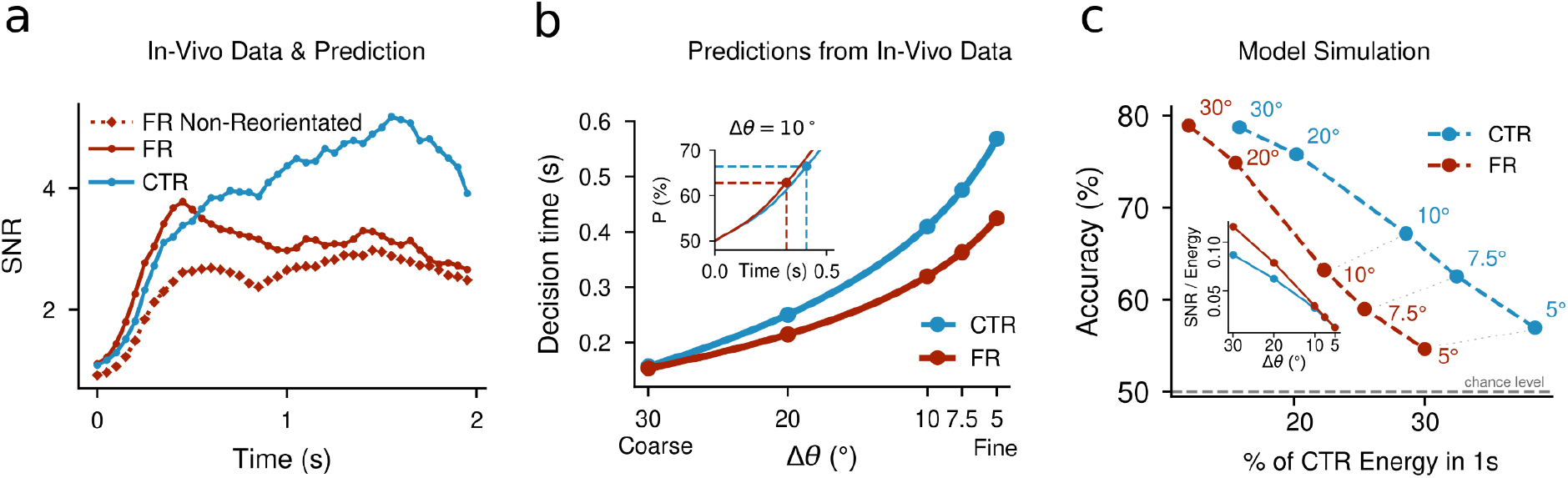
Population coding dynamics predict shorter decision times under food restriction. **a:** SNR calculated per ms time bin of the calcium imaging data. We find that food-restricted SNR increases faster at the beginning, but plateaus at a lower maximum. To illustrate the benefits of the reorientation of the noise as seen in Fig. 1e, we show a putative food-restricted SNR trajectory with the same signal-noise angles as in the control dataset. **b:** We fit the behavioural decision time so that the accumulated SNR in the V1 data matches behavioural performance. We find that finer discrimination tasks take longer, and in general food-restricted animals make decisions faster. **Inset:** The performance, *P*, calculated from the accumulated SNR is plotted against the time since stimulus onset. The decision time is chosen as the time at which *P* is equal to the behavioural performance found in ^1^. **c:** Dynamical CANN model simulation using the input gain inferred from the data shows that for gross discriminations the same performance is met with lower energetic cost; for finer discrimination energy savings are seen together with a decrease in the performance. The energy is defined as the total synaptic current as above and normalised against the energy used in 1s during the control trajectory. **Inset:** Calculating the SNR per unit of Energy shows that food-restricted mice are more energy efficient for gross discriminations. For finer discriminations, the ratio is the same.

In addition, the dynamical trajectory of the SNR offers multiple time points to further validate our linear-response theory. Fitting the relationship between signal-projection ratios and covariance-eigenvalue ratios (see Methods) reproduced the noise covariance well (*R*^2^ = 0. 88) and allowed us to calculate, from calcium-imaging data, an effective food-restricted-to-control current ratio of 0.91, in good agreement with the electrophysiological estimate of 0.87 ^1^; this current ratio returned to one after recovery of the body weight (Extended Data Fig. 8).

We then asked whether the temporal dynamics of SNR could help explain why food-restricted animals showed behavioural performance comparable to controls on a gross visual discrimination task but were impaired on fine discriminations ^1,11^. To do this, we cumulatively integrated the SNR over time and converted it into predicted discrimination performance. For each angular difference, we fit the SNR-derived performance to the observed behavioural performance to estimate the decision time (Fig. 3b Inset and Methods). Figure 3b shows how the estimated decision time scales with task difficulty, consistent with previous studies of decision-making ^12, 13^. In particular, it was found in normally fed mice that optogenetic silencing of V1 from 80 ms after stimulus onset impaired fine (15°) discrimination more strongly than coarse (40°) discrimination ^14^.

Additionally, our fit predicts that food-restricted animals make decisions earlier than control animals, especially for fine discriminations that would require long and metabolically costly integration. Faster decision making was also observed in other species under food restriction, such as Drosophila^15^.

Together, these results suggest that the interaction between SNR dynamics and evidence-integration time preserves gross-discrimination performance in food-restricted animals while selectively impairing their fine-discrimination performance.

### Dynamical simulations uncover performance-energy trade-off

To explore the relationship between SNR and energy dynamics in food-restricted mice, we used the CANN to simulate the SNR trajectory and recorded the energetic costs associated with this, using synaptic currents as a proxy for energy rate. To simulate the dynamics of the SNR, we introduced a time-dependent gain into our model (Eq. 2), previously held fixed between conditions, consistent with recent literature on temporal input dynamics^16^. This term multiplies the external input *I*_*i*_ (θ) and its trajectory ratio was fitted from the calcium-imaging data (see Methods). The fits reveal that food-restricted mice exhibit an initially higher input gain that decays over time, in agreement with a simple argument pointing to high input gain as an energy-efficient strategy under food restriction (Extended Data Fig. 8). We recorded the SNR, fitted it to the behavioural performance in the same way as for Fig. 3b, and then estimated the energy spent in making the particular decision. Results are reported in the energy-accuracy plane in Fig. 3c, where we observe that, for coarse discrimination tasks, the energy consumption is lower in our network model of the food-restricted state compared to controls while maintaining comparable performance. For fine discrimination, the energy consumption is still lower in the food-restricted state due to the short integration time, but performance is lower than in the control state. We see in the Inset of Fig. 3c that the food-restricted performance is more energy efficient for coarse discriminations, as early SNR increases are exploited at reduced cost. Finer discriminations involve longer integration, where the food-restricted evidence advantage diminishes; its reduced accumulated evidence then balances its energy savings. Furthermore, based on the CANN, we ran counterfactual tests on different energy-efficient strategies. We found, in particular, that an alternative scenario of reducing the currents and not increasing the noise resulted in much worse performance (Extended Data Fig. 6b).

## Discussion

Our study sheds light on the population-level, temporal and circuit mechanisms making computations effective at low energy expenditure, as well as their limitations. The increase in variability in the population activity accompanying the decrease in synaptic currents is rendered benign by a reorientation of the dominant noise modes away from directions relevant for discrimination. Our theory reveals that the preservation of SNR arises because signal and noise enter through the same pathway. This provides a general principle by which neural circuits can accommodate increased variability while maintaining reliable computation, and suggests a potential strategy for designing energy-efficient algorithms. The same mechanism could preserve population coding under other perturbations that alter cellular gain or synaptic efficacy^17^, including pharmacological or disease-related changes. The reported reorientation allows food-restricted animals to transiently boost their SNR early after stimulus onset. This early advantage compensates for shorter integration times, preserving coarse discrimination while reducing the overall energetic cost of a decision. Fine discriminations, however, require evidence to be accumulated for a longer time; consequently, earlier decision-making impairs performance, in agreement with behavioural observations for food-restricted mice ^1^. Further behavioural experiments under food restriction would be needed to verify this prediction explicitly.

## Online Methods

### Dataset provenance and experimental animals

All neural recordings and behavioural measurements analysed in this study were originally collected and reported by Padamsey et al.^1^ For the present calcium-imaging reanalysis, the dataset comprised eight food-restricted and six control animals. The animals were adult male C57BL/6J mice. Control mice had *ad libitum* access to food. For food-restricted mice, daily food intake was gradually reduced to achieve a 15% loss of body weight from baseline *ad libitum* body weight. Recordings were performed after 2-3 weeks of food restriction. All mice were weighed and handled daily. To reduce the impact of hunger on recordings, animals were sated with *ad libitum* access to food for approximately 1–3 h before experimental assessment. A subset of food-restricted animals was subsequently returned to *ad libitum* feeding for 2 weeks; recordings obtained after recovery of body weight were treated as the recovery condition. This defines the hungry, sated and recovery sessions. The behavioural datasets were analysed at the animal level. All animal experiments were approved by the University of Edinburgh Animal Welfare and Ethical Review Board and were performed under a UK Home Office project licence in accordance with the Animals (Scientific Procedures) Act 1986.^1^

### Two-photon calcium imaging and visual stimulation

Left V1 was injected with flexed GCaMP6s (AAV1.Syn.Flex.GCaMP6s.WRE.SV40, Addgene) and Cre, under the CamKII promoter (AAV1.CamKII,0.4.Cre.SV40, Addgene), to restrict expression to excitatory neurons. A cranial window was placed over the injected area. Mice were head-fixed under a resonant-scanning two-photon microscope. GCaMP6s was excited at 920 nm and images were acquired at 40 Hz. Imaging was performed in Layer 2/3, approximately 160–280 µm below the cortical surface.

Visual stimuli were generated in MATLAB using Psychtoolbox and presented on an LCD monitor positioned approximately 20 cm from the contralateral eye. Drifting sinusoidal gratings were presented at 12 orientations, spaced by 30° from 0° to 330°. Each grating was presented for 2 s at a temporal frequency of 1 Hz. Grating presentations were separated by 4-s grey-screen intervals, with 4-s black-screen periods at the beginning and end of each trial. Spatial frequencies were 0.02, 0.04, 0.08, 0.16 and 0.32 cycles per degree. Each stimulus condition was repeated across approximately 10 trials in the present reanalysis. Natural-movie stimuli consisted of 60-s video clips repeated approximately 10 times.

The starting data for the present analyses were the fluorescence traces obtained from the published imaging dataset. Motion correction, region-of-interest extraction, neuropil correction and conversion to ΔF/F were performed as described by Padamsey et al.^1^ For the present study, the resulting ΔF/F traces were converted to inferred neural activity using CASCADE^18^. Activity was aligned to stimulus onset and analysed in fixed time bins. Unless otherwise stated, all correlation, population-geometry and inference analyses used the CASCADE-inferred activity rather than raw fluorescence.

For analyses of stimulus-evoked responses, neurons were excluded when their trial-to-trial variability during the grey-screen baseline exceeded their variability during the stimulus epoch. This exclusion criterion was applied before calculating population statistics and was defined independently for each animal and dataset.

### Behavioural task

Behavioural measurements were obtained in separate freely swimming experiments. Mice were trained in a visual discrimination water-maze task in which they learned to associate a target grating orientation with the location of a hidden escape platform. Training began with a relatively easy discrimination between a 90° target grating and a 45° non-target grating. After the animals reached the training criterion, they were tested with progressively smaller orientation differences. The tested angular separations were 45°, 30°, 20°, 10°, 7.5° and 5°, corresponding to target/non-target pairs of 90°/45°, 90°/60°, 90°/70°, 90°/80°, 90°/82.5° and 90°/85°, respectively.

Testing consisted of approximately 10 trials per day for 1–2 days per angular separation, as specified in the original protocol. A 0° separation condition was used to estimate chance performance, and performance on the coarse discrimination was reassessed at the end of testing.

A choice was classified as correct when the mouse selected the side or arm associated with the target orientation and reached the hidden platform; selection of the non-target location was classified as incorrect. Swim time was measured from release into the maze until the mouse reached the platform using video-based tracking.

### Noise correlations and covariance estimation

Noise correlations quantify correlated trial-to-trial fluctuations between neurons during repeated presentations of the same stimulus. Let 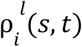 denote the activity of neuron *i* in time bin *t* on trial *l*, in response to stimulus *s*, where each stimulus is defined by a particular angle and spatial frequency. For each stimulus and time bin, we computed

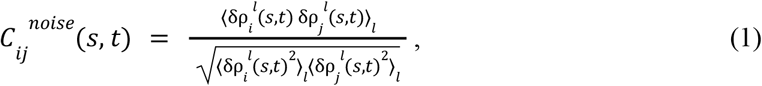

where δρ = ρ − µ is the residual population activity (in vector notation) and µ = ⟨ρ⟩_*l*_ denotes an empirical average over the approximately ten trials of the stimulus *s*. The noise correlation was Fisher-z transformed and averaged across neuron pairs, stimuli and 50-ms time bins. The analysis was performed using the deconvolved calcium signal inferred using CASCADE^18^.

### Population geometry and dimensionality

The noise covariance matrix for each stimulus is given by

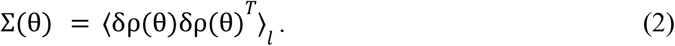

For two nearby stimuli with angles θ and θ + Δθ, the signal vector can be defined as the difference between their population means,

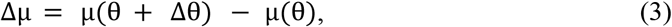

The signal-to-noise ratio can be derived from the linear Fisher information, or equivalently the optimal linearly discriminable boundary^19^ (See Supplementary Information),

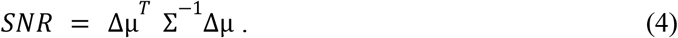

To relate discriminability to the geometry of the noise, we decompose the covariance matrix as Σ *x* = λ *x*, where λ is the noise variance along covariance eigenmode *x*_*k*_. The SNR can then be written as

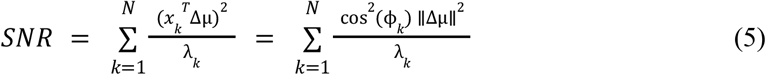

where ϕ_*k*_ is the angle between the signal vector Δµ and the noise mode *x*_*k*_. This recovers Eq. (1) from the main text. Practically, for Fig. 1f, we select one stimulus, calculate the overlap between each noise mode and the closest stimulus and then average over stimuli and animals. The dimensionality was calculated as the participation ratio of each time-resolved covariance spectrum during grating presentation and averaged across 50-ms time bins and stimulus conditions. The participation ratio is defined as

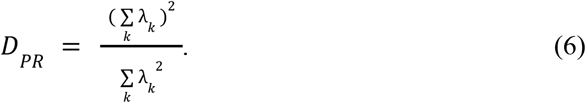

*D*_*k*_ ranges from 1 (if one eigenvalue is non-zero) to *N* (when all eigenvalues are equal).

### Principal-component analysis and signal–noise geometry

To visualise population geometry (Fig. 1b), we applied PCA to the trial-averaged population responses, centred across stimulus orientations. Population means and stimulus-specific noise covariances were projected onto the first two principal components. Noise ellipses were centred on the projected means, with axes determined by the projected covariance and lengths proportional to the corresponding noise standard deviations. Signal vectors connected the population means of nearby stimuli, whereas noise vectors indicated covariance eigenmodes. Quantitative signal–noise overlaps were calculated using the covariance eigenmodes defined above.

### Population decoding

We assessed stimulus discriminability using a linear discriminant analysis^19^ classifier trained to distinguish six orientations from population responses during stimulus presentation (0–2 s). Performance was evaluated using leave-one-trial-out cross-validation and averaged across spatial frequencies within each animal. Trial-shuffled controls were constructed by independently permuting trial order for each neuron within each stimulus condition, preserving marginal response distributions while disrupting across-neuron trial covariance. The decoding comparison is shown in Extended Data Fig. 1c.

### Time-resolved analysis

Population means and noise covariances were calculated separately for each stimulus condition in 50-ms bins aligned to stimulus onset. Signal vectors connected responses to nearby orientations at the same spatial frequency and time bin. We then calculated covariance eigenvalues, signal–noise overlaps, SNR and participation ratios for each bin. For the dimensionality comparison, participation ratios were averaged across stimulus conditions and bins during stimulus presentation to obtain one value per animal. The SNR trajectories describe discriminability within individual bins; temporal accumulation for the behavioural model is described separately.

### Counterfactual signal–noise alignment

To assess the contribution of altered alignment under food restriction, we recalculated the food-restricted SNR using control signal–noise angles. Food-restricted signal magnitudes |Δµ|^2^ and covariance eigenvalues λ_*k*_ were retained, while the squared cosine factors cos^2^(ϕ_*k*_) in the mode-specific SNR were replaced by the corresponding control values in Eq. (5).

### Recovery and alternative stimuli

The same geometric analyses were applied to recovery recordings to assess whether the changes associated with food restriction reversed following restoration of body mass. For spatial-frequency comparisons, signal vectors connected mean responses to different spatial frequencies at a fixed grating orientation. For natural movies, activity was averaged within successive 1-s segments of the 60-s movie. Population means and noise covariances were estimated across repeated presentations of each segment, and signal vectors connected the mean responses of successive segments. These analyses are shown in Extended Data Fig. 2.

### Gaussian–Wishart-process covariance estimation

To assess robustness to covariance estimation with limited trials, we additionally used the framework of Nejatbakhsh et al.^6^. Population means were modelled with Gaussian processes and noise covariance with a low-rank-plus-diagonal Wishart process. Periodic kernels over orientation and squared-exponential kernels over log spatial frequency allowed information to be shared across neighbouring stimulus conditions. Variational inference yielded smooth, positive-definite covariance estimates. We repeated the covariance eigendecomposition and signal–noise alignment analyses using these estimates (Extended Data Fig. 3).

### Linear-response approximation

Using the membrane potential Eq. (2) in the main text we can derive the linear response matrix (see Supplementary Information) as

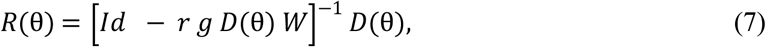

where *Id* is the identity matrix, and *D*(θ) = *F*’[*v*(*t*, θ)] is the effective gain matrix. The neural population response to a small perturbation in stimulus input or presynaptic noise is then

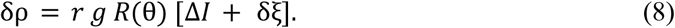

This matrix captures the combined effect of single-cell gain and recurrent amplification. Food restriction changes *R*(θ) through the cellular parameters *r, g*, and *b*, and through the resulting change in the effective gain matrix *D*(θ). As found in ^1^ the input resistance, *r*, increased by 22% and the conductance, *g*, decreased by 29%.

### Transformation of control population responses and tuning curves

We derived in the Supplementary Information the transformation of the eigenvalues of the linear response matrix of the control animals to the food-restricted animals as follows

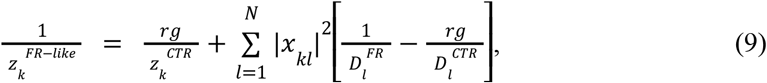

where the linear response decomposition is *R*(θ)*x*_*k*_ = *z*_*k*_ *x*_*k*_, which has the same eigenvectors as the covariance matrix, and *D* is the effective gain matrix. The approximation *D*^*FR*^ = ψ*D*^*CTR*^ was used, where ψ ≈ 1. 24, which was the calculated ratio between the average gains in the data. The agreement between the transformed and experimentally inferred linear-response eigenvalues is shown in Extended Data Fig. 4a.

The transformation was defined as 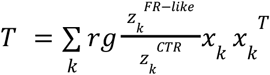, which was applied to both the stimulus means and covariances:

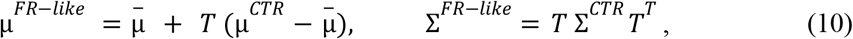

where 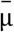 is the midpoint between the two control stimulus means. In Extended Data Fig. 4b, the red ellipses and centres represent the FR reconstruction obtained using the averaged food-restricted linear response eigenvalues, showing excellent agreement with the transformed control responses. To provide the transformation of the tuning curve, we used the following equation

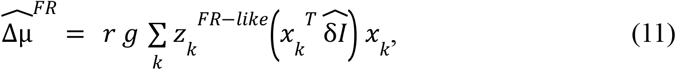

where the input vector 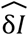 was inferred from the control tuning curves.

### Estimation of effective current ratios and input-gain dynamics

Both the signal vector and covariance ( Eq. (3) and (5) in the main text) depend on the linear response *R*. We can therefore rewrite the ratio between the eigenvalues in terms of the ratio of the signal overlap 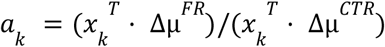, as shown in Supplementary Information, as

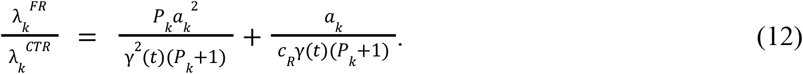

Here, γ(*t*) is the time-dependent ratio between the gains of the input *I*. We model *P*_*k*_ as a time-independent and mode-dependent parameter that depends on the ratio of slow and fast noise components. *c*_*R*_ is the FR/CTR current ratio, which we compare to the electrophysiologically measured quantities denoted in the paper as *r* × *g*. The time-dependent ratio of the gains γ(*t*) was fitted with the inferred *c*_*R*_ in Extended Data Fig. 8c. The current ratio inference was fitted with a time-independent gain ratio γ. Extended Data Fig. 8a,b gives the fit quality and compares the inferred current ratios.

### Stochastic network simulations

To investigate finite-time fluctuations, we simulated microscopic binary dynamics in the same ring network described in ^20^. The state of neuron *i* at simulation step *t* was σ_*i*_ (*t*) ∈ {0, 1}. We simulated a population of *N* = 250 neurons arranged on a one-dimensional ring. Recurrent connectivity was local and symmetric: *W*_*ij*_ = 1/*N* if 0 < *d*_π_ (ϕ_*i*_, ϕ_*j*_) ≤ 0. 05π, and *W*_*ij*_ = 0 otherwise, where *d*_π_ (ϕ_*i*_, ϕ) = *min*(|ϕ_*i*_ − ϕ_*j*_ |, π − |ϕ_*i*_ − ϕ_*j*_ |) is the distance along the ring and the preferred orientation of neuron *i* is ϕ_*i*_ = π(*i* − 1)/*N*. Thus, each neuron received recurrent input from neurons within 5% of the ring in either direction. Self-connections were excluded by setting *W* = 0. A stimulus with centre θ was represented by a Gaussian profile, 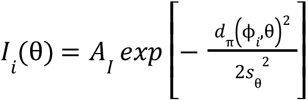, where *A*_*I*_ was the input amplitude and *s*_θ_ its width. Nearby stimuli were generated by shifting the centre θ while keeping the amplitude, width and recurrent connectivity fixed.

Neurons were initialized with *fN* adjacent neurons active at a randomly selected position on the ring. The log probability of the instantaneous neural activity configuration was, up to an irrelevant additive constant,

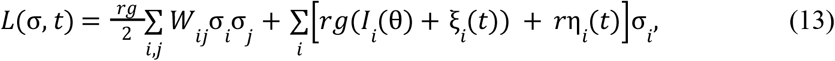

using the noise sources defined above. The fast noise η_*i*_ (*t*) was updated every time step and the slow input noise, ξ_*i*_ (*t*), was updated every 100 time steps. The dynamics were implemented using activity-conserving Metropolis updates. At each update, one active neuron *i* and one inactive neuron *j* were selected randomly, and the exchange (σ_*i*_, σ_*j*_) = (1, 0) → (0, 1) was proposed. The exchange was accepted with probability *P*_*acc*_ = *min*[1, *exp*(Δ*L*)], where Δ*L* is the change in log probability due to the proposed exchange. One simulation sweep consisted of *N* proposed exchanges. In between, around 20% of the time, to allow the total activity to fluctuate, a single neuron flip was proposed. Control and food-restricted simulations used identical connectivity, stimuli and initial activity constraints and differed only in the cellular parameters *g* and *r* defined above.

To examine the effect of temporal averaging, consecutive population states were divided into non-overlapping windows of *K* sweeps and averaged. The mean activity and covariance matrices were calculated across these averaged blocks, and discriminability was quantified using the same signal-to-noise ratio defined for the experimental data. This allowed us to determine how the agreement between the microscopic simulations and the long-time mean-field approximation depended on the averaging window.

The dynamic CANN simulations were performed in the same way, with only the input gain changing over time. The gain ratio followed the time course inferred from the fit and shown in Extended Data Fig. 8, with a brief plateau introduced at its maximum. Motivated by previously reported temporal dynamics of sensory responses^16^, we modelled the trajectory of the control input gain as a steep increasing function until 300 ms after which the slope decreased.

For the alternative scenarios simulated on the CANN, we checked what the performance would look like if the current was reduced to 0.86 without the noise increasing. This is described further in Extended Data Fig. 6b.

### Mean-field theory

Following a standard variational mean-field approach, we represented the mean activity of neuron *i* by the continuous order parameter µ_*i*_ = ⟨σ_*i*_ ⟩. The resulting self-consistency equation was derived in Supplementary Information, following ^20^, to be

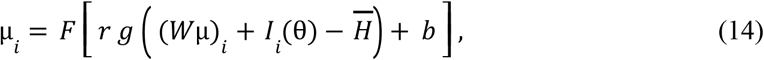

where 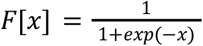 is the logistic transfer function, 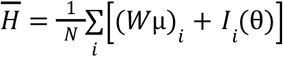 was the global inhibitory current defined to match the population averaged input and *b* was the baseline change. Stable solutions of this equation correspond to local minima of the mean-field free energy. We solved this fixed-point equation using damped updates. To include fast cell-intrinsic fluctuations, the transfer function was averaged over a Gaussian variable:

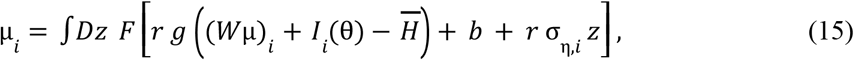

where 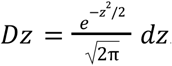. Here, *z* is a standard Gaussian variable and σ_η,*i*_ is the standard deviation of the intrinsic input fluctuations of neuron *i*, which was picked to match the range of the eigenvalues from the data. The Gaussian integral was evaluated numerically using Gauss-Hermite quadrature. After convergence, the effective single-cell gains and the network linear response matrix were evaluated at the resulting operating point. Signal and covariance statistics were then calculated analytically using the linear response expressions described above. The total synaptic input was used as an energy proxy: 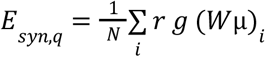.

### Prediction of discrimination performance and inference of integration time

For the behavioural model we calculated the SNR per 50-ms time bin and then cumulatively summed the SNR over time, 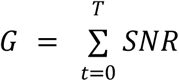. This summation assumes that the neural responses in successive 50-ms bins are statistically independent, so that their contributions to discriminability are additive. We converted the SNR into performance using 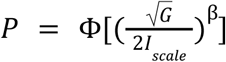 where Φ is the Gaussian cumulative distribution function. We fitted *I*_*scale*_ and selected the β that minimises the mean squared error. Calcium-imaging recordings were available only for 30 degree separations. We simulated the SNR of smaller angular separations by scaling it by sin^2^ (Δθ), proportional to the squared chord length. We then fitted the behavioural decision time so that the performance given by the accumulated SNR matched the behavioural performance from ^1^. Fig. 3b Inset shows how the 10 degree decision time was calculated.

## Acknowledgements

We thank Laurent Bourdieu, Simona Cocco, Danai Katsanevaki and Arno Onken for comments on the manuscript.

R.M. acknowledges support from the French National Research Agency (ANR) through the ENACT project (ANR-25-CE30-0812). Z.P. acknowledges support from the Qatar Foundation. N.L.R. acknowledges support from the Biotechnology and Biological Sciences Research Council (BBSRC; grant BB/T007907/1 to N.L.R. and Z.P.), the Wellcome Trust and the Royal Society (Sir Henry Dale Fellowship), the Shirley Foundation, the Patrick Wild Centre, the RS MacDonald Charitable Trust (Seedcorn Grant), and the Simons Initiative for the Developing Brain. This work received funding from the European Research Council (ERC) under the European Union’s Horizon 2020 research and innovation programme (grant agreement No. 866386).

