## Extended Data for "Neural mechanisms for resilient sensory computation under metabolic stress"

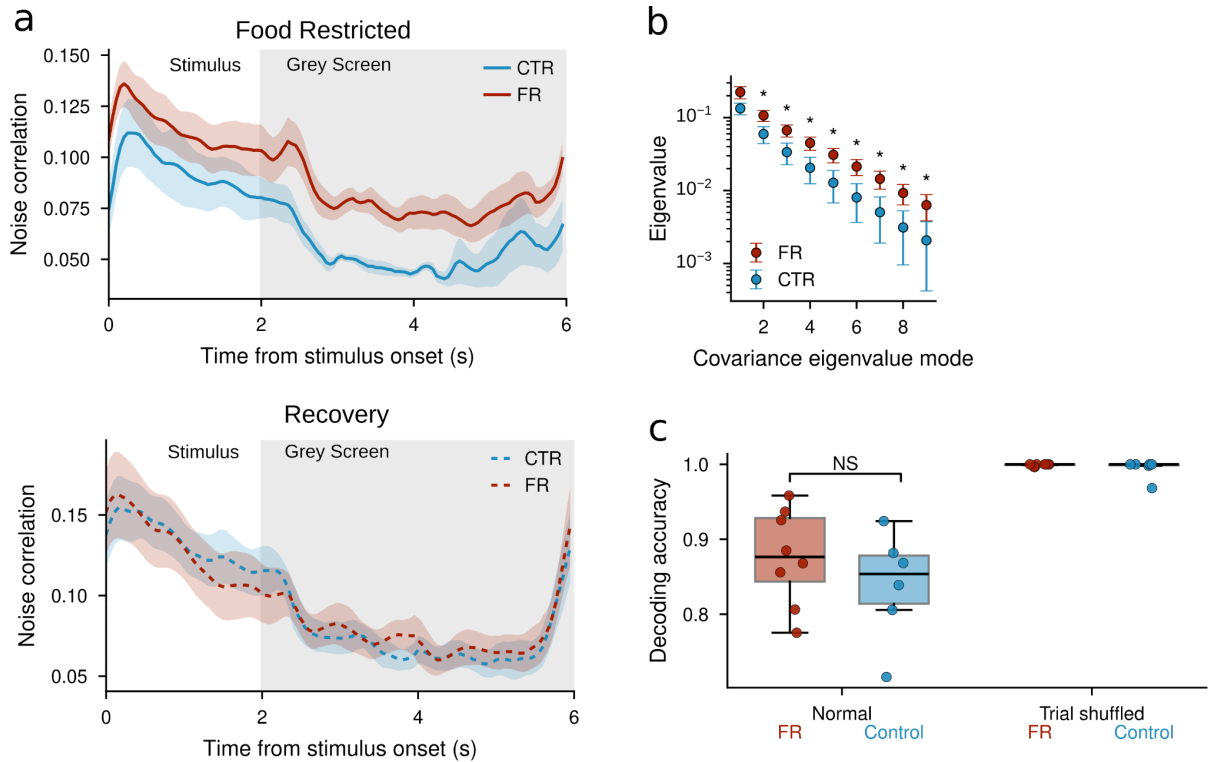

### Extended Data Fig. 1: Noise correlations, covariance spectra and population decoding under food restriction.

**a:** Average time course of noise correlations in food-restricted (body mass reduced by 15% but fed right before the experiment) and recovery (food-restricted animals returned to ad libitum feeding and recorded after 2 weeks of recovery) sessions. Mean noise correlation is shown as a function of time relative to stimulus onset for food-restricted and control animals ($n = 8$ FR and $n = 6$ CTR). Noise correlations were calculated separately at each 50-ms time bin from stimulus-conditioned trial residuals, Fisher-$z$ transformed, averaged within each animal, and back-transformed to $r$ for display. Lines show the mean across animals and shaded regions show the SEM. Grey shading denotes presentation of a grey screen in between stimuli (2–6 s). **b:** Food restriction increased trial-to-trial covariance eigenvalues across multiple population modes. For each stimulus condition and 50-ms time bin during bar presentation (0–2 s), the covariance matrix of deconvolved activity across repeated trials was eigendecomposed. Ordered eigenvalues were averaged across time bins and stimulus conditions within each animal. Points and error bars show the mean ± SEM across animals. Asterisks indicate $p < 0.05$, determined separately for each mode using two-sided permutation tests on log-transformed eigenvalues. **c:** Performance of neural code was preserved despite increased correlations (N.S. $p = 0.142$). Trial-shuffled responses perform perfectly. This supports the hypothesis that noise correlations are in general harmful for discriminability, but the increase in noise correlations arising from food restriction is not. A linear discriminant analysis classifier was trained to decode six orientations from population responses during bar presentation using leave-one-trial-out cross-validation; decoding accuracy was averaged across spatial frequencies within each animal. Trial-shuffled tests to mimic the removal of noise correlations were generated by independently permuting trial order for each neuron, disrupting across-neuron trial covariance while preserving marginal response distributions. Points represent individual animals and boxes show the median and interquartile range. Chance performance was 1/6.

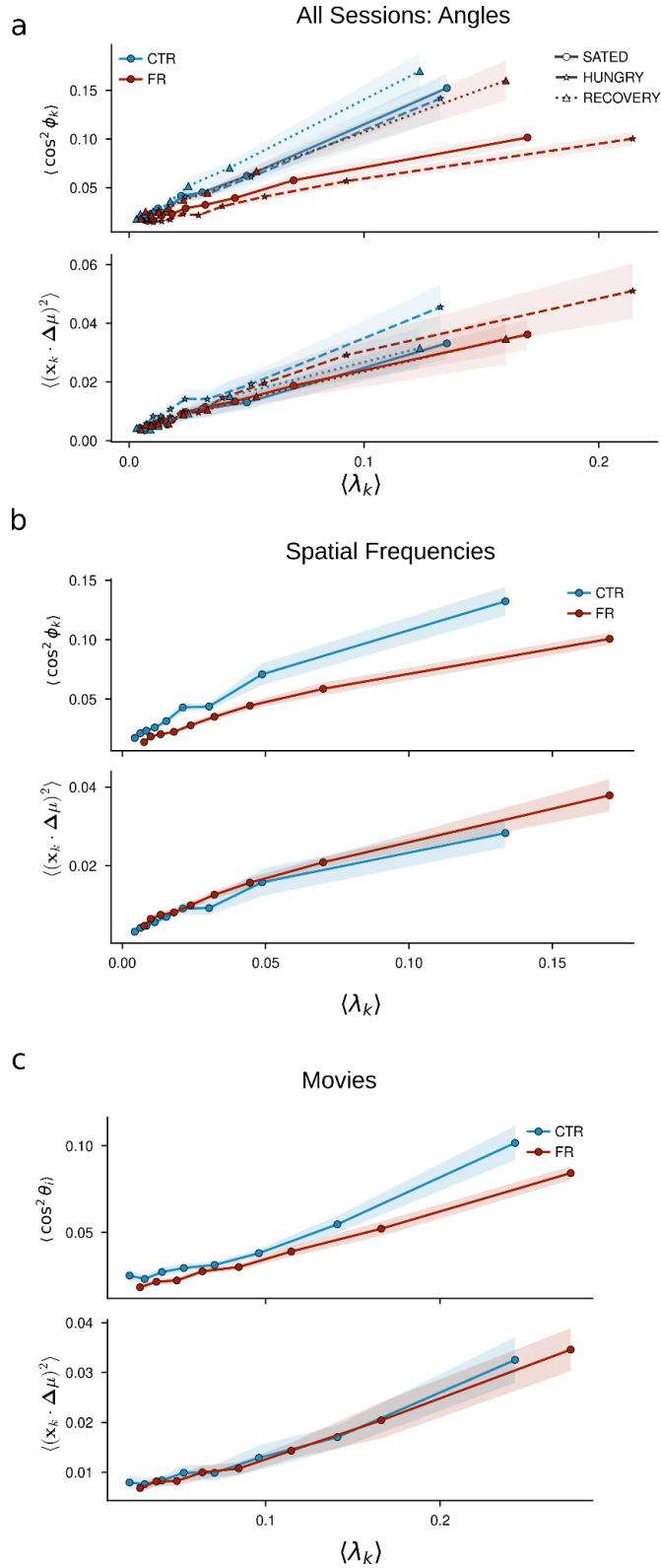

**Extended Data Fig. 2: Signal–noise geometry across recovery sessions and stimulus types.** **a:** Plotting all sessions together (Hungry, Sated and Recovery; see Methods), we see that the squared signal–noise cosine returns to the control baseline after recovery, as in Fig. 1e. Additionally, the overlap–eigenvalue plot is the same for all conditions and sessions. This helps validate the analysis conducted in Fig. 1f. **b:** By defining the response vector between different spatial frequencies of the presented gratings we obtain qualitatively similar results to those in Fig. 1e,f, where the response vector is defined between nearby angles. **c:** By defining the response vector between successive 1s averaged responses from V1 during presentation of a 60-s video, we see qualitatively identical results to Figure 1e and 1f. This highlights the stimulus-independent nature of the food-restricted effect.

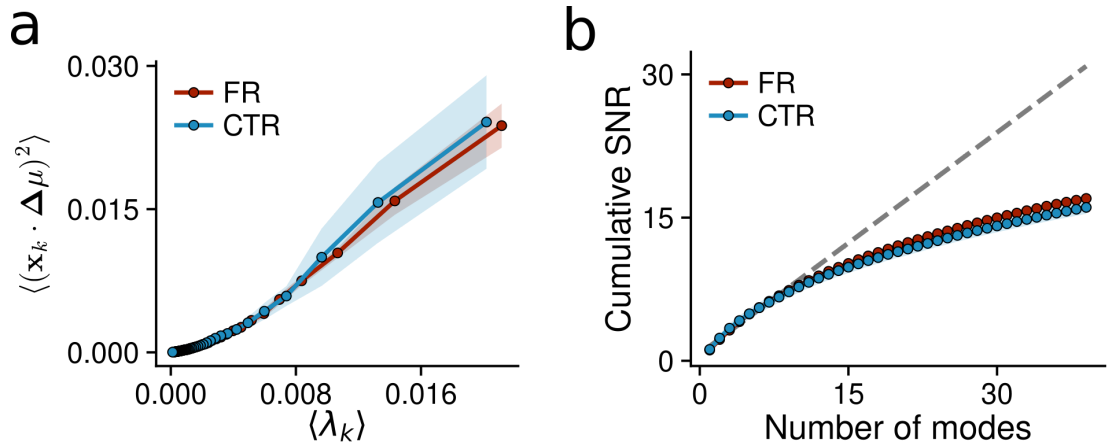

**Extended Data Fig. 3: Gaussian-Wishart-process estimation of stimulus-dependent noise covariance.** Because the number of trials per stimulus condition is small relative to the number of simultaneously recorded neurons, condition-wise empirical covariance matrices are noisy and rank deficient. As a model-based robustness check, we estimated the population mean with Gaussian processes and the full noise covariance with a low-rank-plus-diagonal Wishart process<sup>6</sup>. Periodic and squared-exponential kernels over grating orientation and log spatial frequency, respectively, pool information across nearby stimulus conditions while retaining condition-specific covariance structure. Variational inference therefore provides smooth, positive-definite covariance estimates and access to the complete covariance eigenspectrum. **a:** Replication of the analysis in Fig. 1f using the inferred covariance matrices. The squared overlap is plotted against the corresponding inferred covariance eigenvalue. The overlap-eigenvalue relationship is similar in food-restricted (FR) and control (CTR) animals. **b:** Plotting the cumulative SNR against descending eigenvalue mode, one finds that smaller eigenvalues contribute less to the SNR and begin to saturate. The differences between food-restricted and controls are not statistically significant. This result is for the signal averaged over the 2s exposure to the stimulus. The first 10 modes contribute the same SNR and seem to be roughly linear as found in Fig. 1f. The grey dashed line is a linear fit on the first 10 modes and shows how the SNR would grow if Eq. (4) applied.

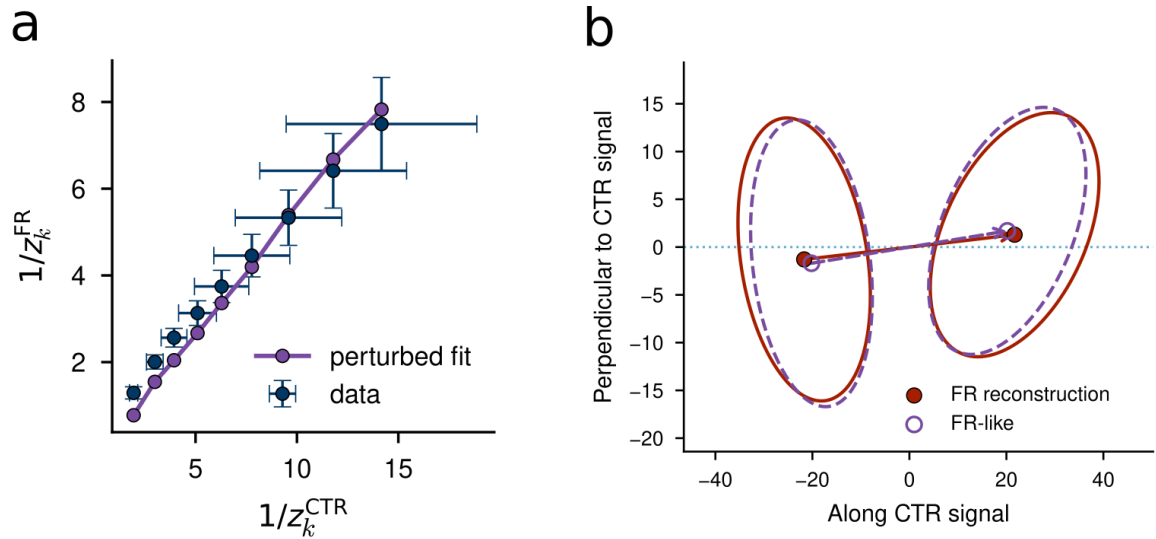

**Extended Data Fig. 4: Transformation of the linear response under food restriction.** **a:** Inverse linear-response eigenvalues from the V1 data in control (CTR) and food-restricted (FR) conditions, compared with the values predicted by the perturbative transformation in Methods Eq. (9) (purple). **b:** Comparison of the FR reconstruction (red), obtained using the measured FR linear-response eigenvalues, with the FR-like responses (purple dashed) predicted by transforming the CTR population means and covariance matrices according to Methods Eq. (10). Points indicate stimulus means and ellipses represent noise covariances. The projection plane is spanned by the CTR and FR signal vectors, with the horizontal axis aligned with the CTR signal and the vertical axis perpendicular to it. The close agreement demonstrates that the transformation reproduces both the changes in signal separation and the geometry of the noise.

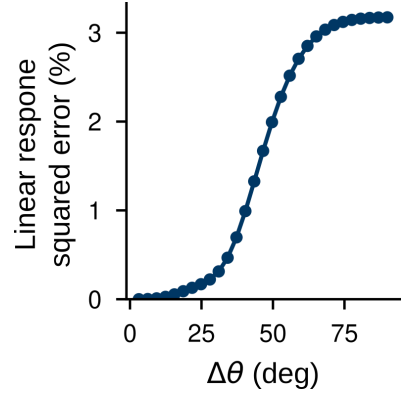

**Extended Data Fig. 5: Accuracy of the linear response approximation across stimulus separations.** From the mean-field analysis detailed in Methods, we produced Fig. 2d,e. The mean-field simulations and theory which led to Eq. (3)-(5) assume that the two stimuli presented are close to each other. This plot shows the error between the mean-field and simulated values of the linear response derived signal vector. The small value of the squared error highlights the validity of the linear response approximations even for a large difference in stimuli. The saturation is seen when the two responses have no overlap.

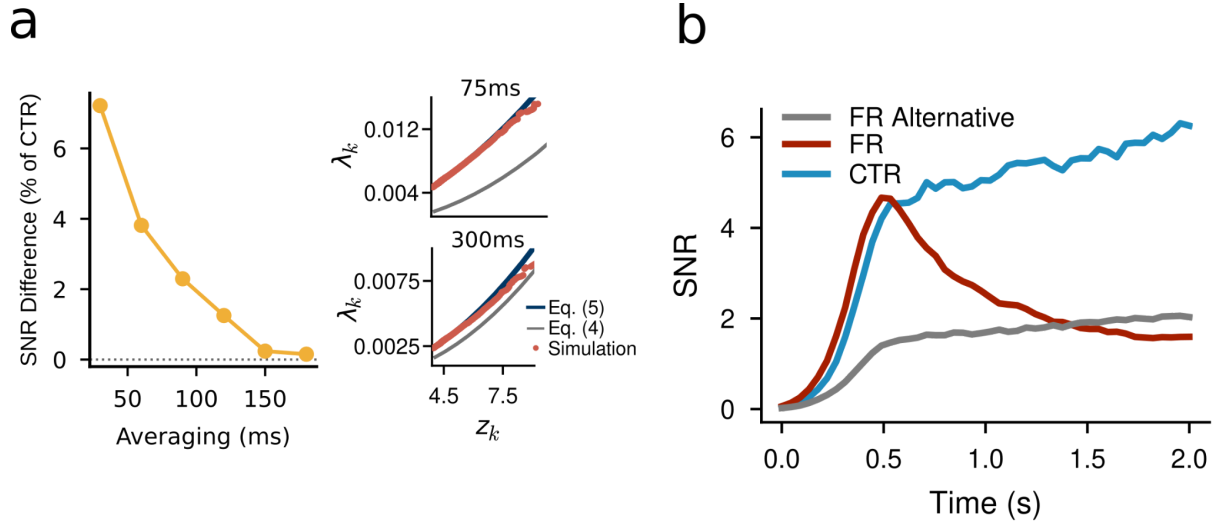

**Extended Data Fig. 6: Finite-time noise fluctuations and alternative energy-saving strategies in circuit simulations.** **a:** Relative difference of SNR, $100 \times (SNR_{CTR} - SNR_{FR})/SNR_{CTR}$, between control and food-restriction vs. averaging window duration in a static CANN simulation. A larger averaging window reduces the linear term in Eq. (5) so that the approximation in the previous model, Eq. (4), holds. **Inset:** relationship between covariance eigenvalues, $\lambda$, and linear response eigenvalues, $z$, for two averaging windows. The panels show how the more complete theory for the covariance, from Eq. (5), reproduces the microscopic simulation and how it coincides with the simpler expression, from Eq. (4), for larger windows. **b:** Population SNR under alternative strategies for reducing energy consumption. In microscopic, dynamical, CANN simulations, we model the dynamics of the SNR over time. For the FR ‘Alternative’ scenario, we maintain the input resistance at 1 but reduce the current to 0.86 in such a way that the total current used is the same between FR and FR Alternative. We see that the SNR of the FR Alternative dynamics is worse than FR, highlighting the computational benefit of the higher noise state.

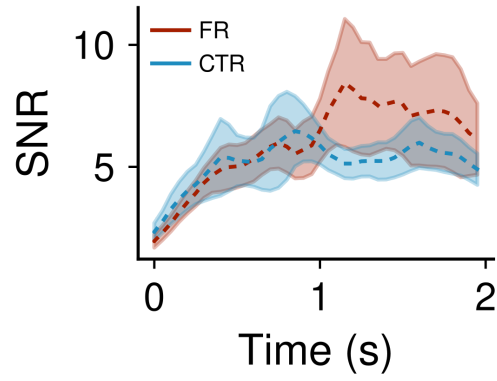

**Extended Data Fig. 7: Population SNR dynamics following recovery from food restriction.** Time-resolved SNR in previously food-restricted (FR, red) and control (CTR, blue) animals during recovery sessions, recorded after 2 weeks of ad libitum feeding and restoration of body mass. SNR was calculated in 50-ms bins relative to stimulus onset using the same procedure as in Fig. 3a. Dashed lines show means across animals and shaded regions indicate SEM. The lower sustained SNR observed during food restriction was no longer apparent after recovery.

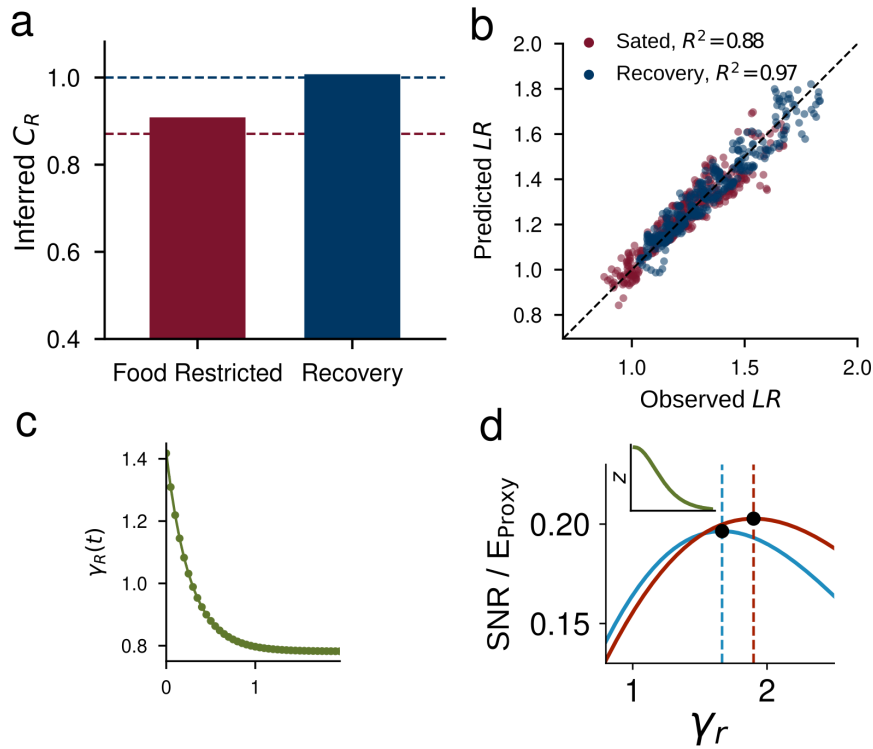

**Extended Data Fig. 8: Inference of the current ratio and input gain.** **a:** Inferred FR/CTR effective current ratio, $c_R$, for each session type with an independent gain ratio as described in Methods. The inferred ratio was 0.909 during the food-restricted (burgundy) condition and returned to approximately unity after recovery (navy), consistent with the experimental reference values indicated by the dashed lines. **b:** Model-predicted versus observed linear-response eigenvalue ratios, $z_{FR}/z_{CTR}$, for both sessions. The black dashed line denotes equality between observed and predicted values. Coefficients of determination were ($R^2=0.88$) for sated sessions and ($R^2=0.97$) for recovery sessions. **c:** Inferring the gain ratio by fitting the noise and overlap ratio from the calcium imaging data. We fixed the inferred current ratios from part **a** and predicted a time dependent external input gain which is larger for food restriction for the first 500 ms. **d:** SNR per unit Energy proxy as a function of gain ratio, $\gamma_r$. SNR was calculated using Eq. (5) from the main text, with linear-response eigenvalues proportional to $1/\cosh^2(c_R \gamma_r/2)$, as illustrated in the inset. The energy proxy was the input gain ratio multiplied by the current ratio. Food restriction (red) shifts the peak to a higher gain ratio, indicating that greater gain maximizes energy efficiency in this condition.
