## Supplementary Information for "Neural mechanisms for resilient sensory computation under metabolic stress"

### 1 Derivation of SNR

#### 1.1 Derivation from Linear Discriminant Analysis

Let $\boldsymbol{\rho} \in \mathbb{R}^N$ denote the population activity of the $N$ neurons in response to a stimulus $\theta$. We consider the discrimination of two nearby stimuli, $\theta$ and $\theta + \Delta\theta$. Under the Gaussian approximation, the population responses are distributed as

$$\begin{aligned}\boldsymbol{\rho} \mid \theta &\sim \mathcal{N}(\boldsymbol{\mu}(\theta), \boldsymbol{\Sigma}), \\ \boldsymbol{\rho} \mid \theta + \Delta\theta &\sim \mathcal{N}(\boldsymbol{\mu}(\theta + \Delta\theta), \boldsymbol{\Sigma}),\end{aligned}\tag{1}$$

where $\boldsymbol{\mu}(\theta) \in \mathbb{R}^N$ is the mean population response and $\boldsymbol{\Sigma} \in \mathbb{R}^{N \times N}$ is the response covariance matrix. Linear Discriminant Analysis, in its simplest formulation, assumes that the covariance is the same for the two stimuli.

The corresponding conditional probability density is

$$p(\boldsymbol{\rho} \mid \theta) = \frac{1}{(2\pi)^{N/2} \det(\boldsymbol{\Sigma})^{1/2}} \exp \left[ -\frac{1}{2} (\boldsymbol{\rho} - \boldsymbol{\mu}(\theta))^\top \boldsymbol{\Sigma}^{-1} (\boldsymbol{\rho} - \boldsymbol{\mu}(\theta)) \right].\tag{2}$$

For equally probable stimuli, the Bayes-optimal classifier assigns the response to $\theta + \Delta\theta$ whenever

$$p(\boldsymbol{\rho} \mid \theta + \Delta\theta) \geq p(\boldsymbol{\rho} \mid \theta).\tag{3}$$

The corresponding log-likelihood ratio is

$$\delta(\boldsymbol{\rho}) = \ln \left[ \frac{p(\boldsymbol{\rho} \mid \theta + \Delta\theta)}{p(\boldsymbol{\rho} \mid \theta)} \right].\tag{4}$$

We define the signal vector as

$$\Delta\boldsymbol{\mu} = \boldsymbol{\mu}(\theta + \Delta\theta) - \boldsymbol{\mu}(\theta).\tag{5}$$

Expanding the two Gaussian likelihoods gives

$$\begin{aligned}\delta(\boldsymbol{\rho}) &= -\frac{1}{2} (\boldsymbol{\rho} - \boldsymbol{\mu}(\theta + \Delta\theta))^\top \boldsymbol{\Sigma}^{-1} (\boldsymbol{\rho} - \boldsymbol{\mu}(\theta + \Delta\theta)) \\ &\quad + \frac{1}{2} (\boldsymbol{\rho} - \boldsymbol{\mu}(\theta))^\top \boldsymbol{\Sigma}^{-1} (\boldsymbol{\rho} - \boldsymbol{\mu}(\theta)) \\ &= \Delta\boldsymbol{\mu}^\top \boldsymbol{\Sigma}^{-1} \boldsymbol{\rho} - \frac{1}{2} \Delta\boldsymbol{\mu}^\top \boldsymbol{\Sigma}^{-1} [\boldsymbol{\mu}(\theta + \Delta\theta) + \boldsymbol{\mu}(\theta)].\end{aligned}\tag{6}$$

The discriminant is therefore linear in the population response,

$$\delta(\boldsymbol{\rho}) = \mathbf{w}^\top \boldsymbol{\rho} + b_{\text{LDA}},\tag{7}$$

where

$$\begin{aligned}\mathbf{w} &= \Sigma^{-1} \Delta \boldsymbol{\mu}, \\ b_{\text{LDA}} &= -\frac{1}{2} \mathbf{w}^\top [\boldsymbol{\mu}(\theta + \Delta\theta) + \boldsymbol{\mu}(\theta)].\end{aligned}\tag{8}$$

Thus, LDA projects the population response onto

$$y = \mathbf{w}^\top \boldsymbol{\rho}, \quad \mathbf{w} = \Sigma^{-1} \Delta \boldsymbol{\mu}.\tag{9}$$

The difference between the projected means is

$$\Delta \mu_y = \mathbf{w}^\top \Delta \boldsymbol{\mu},\tag{10}$$

and the variance of the projected response is

$$\sigma_y^2 = \mathbf{w}^\top \Sigma \mathbf{w}.\tag{11}$$

The squared signal-to-noise ratio is therefore

$$\text{SNR} = \frac{(\mathbf{w}^\top \Delta \boldsymbol{\mu})^2}{\mathbf{w}^\top \Sigma \mathbf{w}}.\tag{12}$$

Substituting the LDA direction $\mathbf{w} = \Sigma^{-1} \Delta \boldsymbol{\mu}$ gives

$$\begin{aligned}\mathbf{w}^\top \Delta \boldsymbol{\mu} &= \Delta \boldsymbol{\mu}^\top \Sigma^{-1} \Delta \boldsymbol{\mu}, \\ \mathbf{w}^\top \Sigma \mathbf{w} &= \Delta \boldsymbol{\mu}^\top \Sigma^{-1} \Delta \boldsymbol{\mu}.\end{aligned}\tag{13}$$

Hence,

$$\text{SNR} = \Delta \boldsymbol{\mu}^\top \Sigma^{-1} \Delta \boldsymbol{\mu}.\tag{14}$$

### 1.2 Derivation from Linear Fisher Information

We now relate the squared signal-to-noise ratio to the linear Fisher information. Consider a continuously varying stimulus $\theta$ and a population response distributed as

$$\boldsymbol{\rho} \mid \theta \sim \mathcal{N}(\boldsymbol{\mu}(\theta), \Sigma).\tag{15}$$

We assume both the mean response and covariance may depend on the stimulus. We write

$$\boldsymbol{\mu}'(\theta) = \frac{\partial \boldsymbol{\mu}(\theta)}{\partial \theta}, \quad \Sigma'(\theta) = \frac{\partial \Sigma(\theta)}{\partial \theta}.\tag{16}$$

The Fisher information is defined as the variance of the score,

$$J(\theta) = \left\langle \left[ \frac{\partial}{\partial \theta} \ln p(\boldsymbol{\rho} \mid \theta) \right]^2 \right\rangle_{\boldsymbol{\rho} \mid \theta}.\tag{17}$$

Differentiating the Gaussian log-likelihood gives

$$\begin{aligned}\frac{\partial}{\partial \theta} \ln p(\boldsymbol{\rho} \mid \theta) &= \boldsymbol{\mu}'^\top \Sigma^{-1} (\boldsymbol{\rho} - \boldsymbol{\mu}) \\ &\quad + \frac{1}{2} (\boldsymbol{\rho} - \boldsymbol{\mu})^\top \Sigma^{-1} \Sigma' \Sigma^{-1} (\boldsymbol{\rho} - \boldsymbol{\mu}) \\ &\quad - \frac{1}{2} \text{Tr}(\Sigma^{-1} \Sigma'),\end{aligned}\tag{18}$$

where the dependence on $\theta$ is left implicit on the right-hand side.

For Gaussian responses, the Fisher information separates into a contribution from changes in the mean response and a contribution from changes in the covariance,

$$J(\theta) = J_{\text{lin}}(\theta) + J_{\text{cov}}(\theta), \quad (19)$$

where

$$J_{\text{lin}}(\theta) = \boldsymbol{\mu}'(\theta)^\top \boldsymbol{\Sigma}(\theta)^{-1} \boldsymbol{\mu}'(\theta), \quad (20)$$

and

$$J_{\text{cov}}(\theta) = \frac{1}{2} \text{Tr} \left[ \boldsymbol{\Sigma}(\theta)^{-1} \boldsymbol{\Sigma}'(\theta) \boldsymbol{\Sigma}(\theta)^{-1} \boldsymbol{\Sigma}'(\theta) \right]. \quad (21)$$

The first term measures information carried by changes in the mean population response. The second measures additional information carried by stimulus-dependent changes in the covariance. The linear Fisher information is an equivalent measure to the optimal LDA signal to noise ratio (SNR), in the small angle limit: $\text{SNR} = J_{\text{lin}} \Delta \theta^2$.

### 2 Derivation of the Linear Response Theory

We model the population response to an oriented stimulus using a recurrent network of $N$ neurons. The activity of neuron $i$ is written as

$$\rho_i(t) = F[v_i(t)], \quad (22)$$

where $F$ is a transfer function. The membrane potential is

$$v_i(t) = rg \left( \sum_{j=1}^N W_{ij} \rho_j(t) + I_i(\theta) + \xi_i(t) \right) + r\eta_i(t) + b. \quad (23)$$

Here $W_{ij}$ is the recurrent coupling from neuron $j$ to neuron $i$, $I_i(\theta)$ is the external input evoked by stimulus $\theta$, $\xi_i(t)$ is a trial-to-trial presynaptic fluctuation, $\eta_i(t)$ is the cell-intrinsic noise, $r$ is the input resistance, $g$ is the AMPAR-mediated synaptic conductance, and $b$ is an effective threshold or baseline term. The low-power mode induced by food restriction is represented by changing the cellular parameters $r, g, b$, while keeping the recurrent connectivity $\mathbf{W}$, the stimulus input $\mathbf{I}(\theta)$, and the statistics of the noise sources fixed. In Padamsey et al. 2022<sup>1</sup>, these cellular changes explain how neural variability can increase while mean firing rates remain stable.

In the following analysis, we consider the regime in which the measured covariance is dominated by the presynaptic fluctuations $\xi_i(t)$. The fast intrinsic fluctuations $\eta_i(t)$ are incorporated through their effect on the neuronal susceptibility. We capture this effect by defining a noise-averaged transfer function,

$$\rho_i = F_\eta(u) = \langle F(u + r\eta) \rangle_\eta, \quad (24)$$

where

$$u_i(t) = rg \left( \sum_{j=1}^N W_{ij} \rho_j(t) + I_i(\theta) + \xi_i(t) \right) + b. \quad (25)$$

The linear response is then computed with respect to $F_\eta$. From this point onward, $\boldsymbol{\rho}$ denotes the response averaged over the fast intrinsic noise. For a fixed stimulus $\theta$, the mean response is

$$\boldsymbol{\mu}(\theta) = \langle \boldsymbol{\rho}(\theta) \rangle. \quad (26)$$

We consider small perturbations around this mean response,

$$\begin{aligned}\delta\boldsymbol{\rho} &= \boldsymbol{\rho} - \boldsymbol{\mu}, \\ \delta\mathbf{u} &= \mathbf{u} - \langle\mathbf{u}\rangle.\end{aligned}\tag{27}$$

Linearising the effective transfer function gives

$$\delta\boldsymbol{\rho} = \mathbf{D}(\theta) \delta\mathbf{u},\tag{28}$$

where $\mathbf{D}(\theta)$ is a diagonal effective gain matrix,

$$[\mathbf{D}(\theta)]_{ij} = F'_\eta(\langle u_i \rangle) \delta_{ij}.\tag{29}$$

Thus, $\mathbf{D}(\theta)$ is the effective susceptibility of the population at the stimulus-dependent operating point. It measures how small changes in the recurrent or external input are converted into changes in firing activity, after averaging over the fast intrinsic fluctuations. The perturbation to the slow input is

$$\delta\mathbf{u} = rg\mathbf{W} \delta\boldsymbol{\rho} + rg\delta\mathbf{I} + rg\delta\boldsymbol{\xi}.\tag{30}$$

Combining this with $\delta\boldsymbol{\rho} = \mathbf{D}(\theta)\delta\mathbf{u}$ gives

$$\delta\boldsymbol{\rho} = \mathbf{D}(\theta) [rg\mathbf{W} \delta\boldsymbol{\rho} + rg\delta\mathbf{I} + rg\delta\boldsymbol{\xi}].\tag{31}$$

Rearranging,

$$[\mathbf{I}_N - rg\mathbf{D}(\theta)\mathbf{W}] \delta\boldsymbol{\rho} = rg\mathbf{D}(\theta) [\delta\mathbf{I} + \delta\boldsymbol{\xi}].\tag{32}$$

We therefore define the effective local response matrix

$$\mathbf{R}(\theta) = [\mathbf{I}_N - rg\mathbf{D}(\theta)\mathbf{W}]^{-1} \mathbf{D}(\theta),\tag{33}$$

so that

$$\delta\boldsymbol{\rho} = rg\mathbf{R}(\theta) [\delta\mathbf{I} + \delta\boldsymbol{\xi}].\tag{34}$$

This matrix is the central object of the theory. It describes how a small perturbation in the effective input is transformed by both the single-cell gain and the recurrent network. Food restriction changes $\mathbf{R}(\theta)$ through the cellular parameters $r$, $g$, $b$, and through the effective gain $\mathbf{D}(\theta)$, even when the recurrent matrix $\mathbf{W}$ itself is unchanged.

### 2.1 Signal Vector

We next compute the signal vector between two nearby stimuli, $\theta$ and $\theta + \Delta\theta$. Their difference in external input is

$$\Delta\mathbf{I} = \mathbf{I}(\theta + \Delta\theta) - \mathbf{I}(\theta).\tag{35}$$

The signal vector is the difference between the mean population responses,

$$\Delta\boldsymbol{\mu} = \boldsymbol{\mu}(\theta + \Delta\theta) - \boldsymbol{\mu}(\theta).\tag{36}$$

To obtain this from the linearised model, we set the noise perturbation to zero and keep only the stimulus perturbation. This gives

$$\Delta\boldsymbol{\mu} \simeq rg \mathbf{R}(\theta) \Delta\mathbf{I},\tag{37}$$

where we see that the stimulus signal is the input difference after being transformed by the same effective recurrent response matrix $\mathbf{R}(\theta)$.

### 2.2 Covariance

We now compute the covariance due to presynaptic fluctuations. At fixed stimulus, the noise-induced response fluctuation is

$$\delta \boldsymbol{\rho}_{\text{noise}}(t) \simeq rg \mathbf{R}(\theta) \delta \boldsymbol{\xi}(t). \quad (38)$$

Assume that the presynaptic fluctuations have covariance

$$\langle \delta \boldsymbol{\xi} \delta \boldsymbol{\xi}^\top \rangle = \sigma_\xi^2 \mathbf{I}_N. \quad (39)$$

Then the covariance of the population response is

$$\boldsymbol{\Sigma} = \langle \delta \boldsymbol{\rho}_{\text{noise}} \delta \boldsymbol{\rho}_{\text{noise}}^\top \rangle \simeq (rg\sigma_\xi)^2 \mathbf{R}(\theta) \mathbf{R}(\theta)^\top. \quad (40)$$

Both the signal vector and the dominant noise covariance are shaped by the same effective network response. The signal scales as $rg \mathbf{R}(\theta)$, while the covariance scales as $(rg)^2 \mathbf{R}(\theta) \mathbf{R}(\theta)^\top$.

In principle, the intrinsic fluctuations also generate an additional covariance term. However, this contribution is reduced by temporal averaging and is assumed to be subleading compared with the covariance generated by the slower presynaptic fluctuations. We therefore focus on the covariance induced by $\boldsymbol{\xi}$, while retaining the effect of $\boldsymbol{\eta}$ through the effective gain matrix $\mathbf{D}$.

### 2.3 SNR

The discriminability of two nearby stimuli is quantified by the squared signal-to-noise ratio,

$$\text{SNR} = \Delta \boldsymbol{\mu}^\top \boldsymbol{\Sigma}^{-1} \Delta \boldsymbol{\mu}. \quad (41)$$

Substituting the expressions above gives

$$\text{SNR} \simeq \frac{1}{\sigma_\xi^2} \|\Delta \mathbf{I}\|^2. \quad (42)$$

The factors $rg$ and $\mathbf{R}(\theta)$ cancel because the stimulus signal and the presynaptic noise pass through the same effective network filter. Therefore, changes in input resistance, synaptic conductance, recurrent amplification, and effective single-cell gain can strongly reshape the geometry of the population response, increasing both the size and orientation of noise modes, without necessarily changing the total discriminability.

This cancellation can also be written mode by mode. Suppose that $\mathbf{R}(\theta)$ is symmetric, with eigenvectors $\mathbf{x}_k$ and eigenvalues $z_k$,

$$\mathbf{R}(\theta) \mathbf{x}_k = z_k \mathbf{x}_k. \quad (43)$$

For independent presynaptic noise, the covariance eigenvalue along mode $k$ is

$$\lambda_k = (rg\sigma_\xi)^2 z_k^2. \quad (44)$$

The projection of the signal onto the same mode is

$$\mathbf{x}_k^\top \Delta \boldsymbol{\mu} = rg z_k \mathbf{x}_k^\top \Delta \mathbf{I}. \quad (45)$$

Therefore, the contribution of mode $k$ to discriminability is

$$\frac{(\mathbf{x}_k^\top \Delta \boldsymbol{\mu})^2}{\lambda_k} = \frac{1}{\sigma_\xi^2} (\mathbf{x}_k^\top \Delta \mathbf{I})^2. \quad (46)$$

The response matrix eigenvalue $z_k$ cancels mode by mode. This gives a simple geometric interpretation of the experimental result. The network can appear noisier while preserving the discriminability of the population code.

#### 3 Full finite-window covariance and SNR

##### 3.1 Derivation of Complete Noise Covariance

We now compute the covariance of the response averaged over a finite time window. We denote by $L$ the ratio between the duration of this window and the correlation time of the fast cellular noise $\eta$. Let

$$\bar{\rho} = \frac{1}{L} \sum_{t=1}^L \rho(t) \quad (47)$$

be the activity averaged over a response window of length $L$, and $t$ denotes the index of the time interval. Fast noises in two distinct time intervals are by construction uncorrelated. The mean averaged response is

$$\mu = \langle \bar{\rho} \rangle, \quad (48)$$

and the covariance of the averaged response is

$$\Sigma = \langle (\bar{\rho} - \mu)(\bar{\rho} - \mu)^\top \rangle. \quad (49)$$

The natural separation between fast and slow contributions is obtained by conditioning on the slow presynaptic noise trajectory

$$\Xi = \{\xi(1), \dots, \xi(L)\}. \quad (50)$$

For a fixed, slow-noise trajectory $\Xi$, the mean response after averaging over the fast noise is $\mathbf{m}_\Xi(t) = \langle \rho(t|\Xi) \rangle_\eta$. Using the law of total covariance,

$$\Sigma = \langle (\bar{\rho} - \bar{\mathbf{m}}_\Xi)(\bar{\rho} - \bar{\mathbf{m}}_\Xi)^\top \rangle_{\eta, \Xi} + \langle (\bar{\mathbf{m}}_\Xi - \mu)(\bar{\mathbf{m}}_\Xi - \mu)^\top \rangle_\Xi = \langle \Sigma_{\text{Fast}}(\Xi) \rangle_\Xi + \Sigma_{\text{Slow}}, \quad (51)$$

where the $\bar{\mathbf{m}}$ is averaged over time. This decomposition gives a split between fast and slow contributions to the covariance. The first term is the covariance of fast fluctuations within a fixed realisation of the slow presynaptic drive. This gives the fluctuation–dissipation-like contribution as shown below. The second term is the covariance of the conditional mean response as the slow presynaptic drive changes. This gives the linear-response contribution computed above.

###### 3.1.1 Within-state fast term

At fixed $\Xi$, the fluctuations around the conditional mean are $\delta \rho(t) = \rho(t) - \mathbf{m}_\Xi$. Assuming the fast fluctuations are uncorrelated between time bins we can define

$$\Sigma_{\text{Fast}}(\Xi) = \frac{1}{L^2} \sum_{t=1}^L \sum_{t'=1}^L \langle \delta \rho(t) \delta \rho(t')^\top | \Xi \rangle_\eta, \quad (52)$$

To estimate the instantaneous fast covariance, we assume that, conditional on the slow presynaptic drive, the fast variables are described by an exponential distribution

$$P(\rho | \xi) = \frac{1}{Z_\xi} \exp[-H_\xi(\rho)]. \quad (53)$$

This assumption is inspired by equilibrium Gibbs distribution in statistical physics and is correct (at fixed stimulus $\theta$) for the 0-1 neural, Ising-like model simulated in the present work, see Methods. Introducing a small field $\mathbf{f}$ conjugate to the activity,

$$H_\xi(\rho) \longrightarrow H_\xi(\rho) - \mathbf{f}^\top \rho, \quad (54)$$

the susceptibility is by the fluctuation–dissipation relation

$$\chi_{ij} = \frac{\partial \langle \rho_i \rangle_{\eta|\xi}}{\partial f_j} = \langle \delta \rho_i \delta \rho_j \rangle_{\eta|\xi}. \quad (55)$$

Identifying the susceptibility with the linear response matrix derived above, evaluated at the operating point, the within-state fast contribution to the covariance of the averaged response is

$$\mathbf{\Sigma}^{\text{fast}} = \frac{1}{L} \langle \chi(\xi) \rangle_{\xi} \approx \frac{\mathbf{R}}{L}. \quad (56)$$

This is the Ising-like term. Its factor $1/L$ comes directly from averaging fast decorrelated fluctuations over the response window.

#### 3.1.2 Between-state slow term

We now consider the second term, $\mathbf{\Sigma}_{\text{Slow}}$. This term measures how the fast-averaged response changes as the slow presynaptic drive changes. For small slow-noise fluctuations, the linear response result derived above gives

$$\mathbf{m}_{\Xi}(t) - \boldsymbol{\mu} \simeq r g \mathbf{R} \delta \boldsymbol{\xi}(t). \quad (57)$$

Therefore

$$\mathbf{\Sigma}^{\text{slow}} \approx \frac{r^2 g^2}{L^2} \sum_{t,t'=1}^L \mathbf{R} \langle \delta \boldsymbol{\xi}(t) \delta \boldsymbol{\xi}(t')^{\top} \rangle_{\Xi} \mathbf{R}^{\top}. \quad (58)$$

We assume that the slow presynaptic noise covariance show temporal correlations such that if the response window is short compared to the slow noise correlation time:

$$\mathbf{\Sigma}^{\text{slow}} = r^2 g^2 \sigma_{\xi}^2 \mathbf{R} \mathbf{R}^{\top}. \quad (59)$$

This term is therefore not averaged down as $1/L$.

#### 3.1.3 Total covariance

Combining the within-state fast term and the between-state slow term gives

$$\boxed{\mathbf{\Sigma} \simeq \frac{1}{L} \mathbf{R} + r^2 g^2 \sigma_{\xi}^2 \mathbf{R} \mathbf{R}^{\top}}. \quad (60)$$

The first term is the within-state Ising-like covariance of the fast variables, reduced by averaging over $L$ decorrelated samples. The second term is the between-state covariance induced by slow presynaptic fluctuations and obtained by linear response.

### 3.2 Complete SNR Equation

The eigendecomposition of the linear response is $\mathbf{R} \mathbf{x}_k = z_k \mathbf{x}_k$. The eigendecomposition of the covariance is $\mathbf{\Sigma} \mathbf{x}_k = \lambda_k \mathbf{x}_k$. The relation between signal and noise is quantified by the local discriminability

$$\text{SNR} = \Delta \boldsymbol{\mu}^{\top} \mathbf{\Sigma}^{-1} \Delta \boldsymbol{\mu}. \quad (61)$$

If the coupling matrix is symmetric, the squared overlap between the eigenvectors of the noise covariance and the signal vector is defined as

$$\begin{aligned} \text{SNR} &= \sum_k \frac{\cos^2(\phi_k) \|\Delta\boldsymbol{\mu}\|^2}{\lambda_k} \\ &= \sum_k \frac{(\mathbf{x}_k^\top \Delta\boldsymbol{\mu})^2}{\lambda_k} \\ &= \sum_k \frac{(rgz_k)^2 (\mathbf{x}_k^\top \Delta\mathbf{I})^2}{(rg\sigma_\xi z_k)^2 + \frac{z_k}{L}} \\ &= \sum_k \frac{(\mathbf{x}_k^\top \Delta\mathbf{I})^2}{\sigma_\xi^2 + ((rg)^2 L z_k)^{-1}}, \end{aligned} \tag{62}$$

where the overlap $(\mathbf{x}_k^\top \Delta\mathbf{I})$ is taken to be independent of $k$. This is supported by the empirical distributions of the normalized signal projections $(\mathbf{x}_k^\top \Delta\boldsymbol{\mu})/\sqrt{\lambda_k}$. These distributions are approximately Gaussian and vary only weakly across the leading modes (Supplementary Fig. 1), consistent with the approximately linear relationship between squared signal overlap and covariance eigenvalue observed in Fig. 1f.

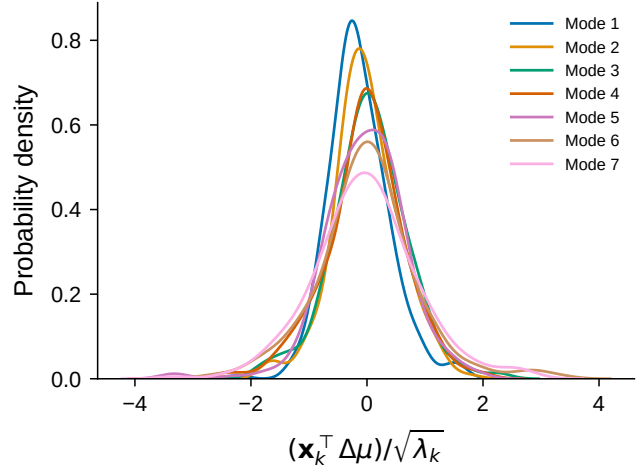

**Supplementary Fig. 1** | Distribution of normalized signal projections across covariance modes. Empirical distributions of $(\mathbf{x}_k^\top \Delta\boldsymbol{\mu})/\sqrt{\lambda_k}$ for the first seven modes for control animals. The distributions show approximately Gaussian shapes and relatively weak dependence on mode index. There was no difference between control and food-restricted.

### 4 Changes to the linear-response spectrum under food restriction

Food restriction changes the response geometry through two related effects. At the single-neuron level, it changes the effective input-output relation of each cell. At the population level, the recurrent network converts the altered single-neuron gains into a global change in the linear-response modes. The relevant single-cell quantity is

$$D_i = \frac{\partial}{\partial u_i} \langle F(u_i + r\eta_i) \rangle_\eta \simeq \langle F'(u_i + r\eta_i) \rangle_\eta. \tag{63}$$

For a sigmoidal transfer function,

$$F'(v) = F(v) [1 - F(v)], \quad (64)$$

so the effective gain is largest near threshold and small in the low-activity and saturated regimes. Food restriction affects $D_i$ in two ways. First, the reduced distance to threshold shifts weakly driven neurons closer to the high-gain region of the input-output curve, shifting the curve to the left. Second, fast intrinsic fluctuations broaden the effective input-output relation, so that neurons sample a greater range of membrane potentials around their mean input. This reduces the peak slope of the input-output curve but extends the region of non-negligible susceptibility. As a result, a neuron $i$ on the shoulders of the stimulus-evoked response can experience a substantial increase to $D_i$.

The recurrent network enters through the response matrix, Eq. (33). Changes in the effective gain $\mathbf{D}$ therefore do not only rescale individual neuronal responses, but also alter how perturbations are propagated through the recurrent circuit. Food restriction shifts the operating point of the network, and modes with stronger recurrent support can be amplified more strongly.

##### 4.1 Mapping the control response matrix to food restriction

For condition $q \in \{\text{CTR}, \text{FR}\}$, the linear response matrix is

$$\mathbf{R}_q(\theta) = [\mathbf{I}_N - (rg)_q \mathbf{D}_q(\theta) \mathbf{W}]^{-1} \mathbf{D}_q(\theta). \quad (65)$$

Taking the inverse gives

$$\mathbf{R}_q^{-1} = \mathbf{D}_q^{-1} - (rg)_q \mathbf{W}. \quad (66)$$

We absorb the control value of the effective recurrent coupling into the definition of the connectivity matrix,

$$\widetilde{\mathbf{W}} = (rg)_{\text{CTR}} \mathbf{W}, \quad (67)$$

and define the relative coupling

$$c_R = \frac{(rg)_{\text{FR}}}{(rg)_{\text{CTR}}} = g_R r_R, \quad (68)$$

where

$$r_R = \frac{r_{\text{FR}}}{r_{\text{CTR}}}, \quad g_R = \frac{g_{\text{FR}}}{g_{\text{CTR}}}. \quad (69)$$

In our parametrisation of the results in [1],

$$c_R \simeq 0.87. \quad (70)$$

The response matrices in the two conditions therefore satisfy

$$\mathbf{R}_{\text{CTR}}^{-1} = \mathbf{D}_{\text{CTR}}^{-1} - \widetilde{\mathbf{W}}, \quad (71)$$

and

$$\mathbf{R}_{\text{FR}}^{-1} = \mathbf{D}_{\text{FR}}^{-1} - c_R \widetilde{\mathbf{W}}. \quad (72)$$

Rearranging Eq. (71) gives

$$\widetilde{\mathbf{W}} = \mathbf{D}_{\text{CTR}}^{-1} - \mathbf{R}_{\text{CTR}}^{-1}. \quad (73)$$

Substituting this expression into Eq. (72) yields

$$\mathbf{R}_{\text{FR}}^{-1} = \mathbf{D}_{\text{FR}}^{-1} - c_R \left( \mathbf{D}_{\text{CTR}}^{-1} - \mathbf{R}_{\text{CTR}}^{-1} \right) \quad (74)$$

$$= \mathbf{D}_{\text{FR}}^{-1} - c_R \mathbf{D}_{\text{CTR}}^{-1} + c_R \mathbf{R}_{\text{CTR}}^{-1}. \quad (75)$$

We can therefore write the FR inverse response matrix as a scaled control inverse response matrix plus a perturbation:

$$\mathbf{R}_{\text{FR}}^{-1} = c_R \mathbf{R}_{\text{CTR}}^{-1} + \mathbf{M}, \quad (76)$$

where

$$\mathbf{M} = \mathbf{D}_{\text{FR}}^{-1} - c_R \mathbf{D}_{\text{CTR}}^{-1}. \quad (77)$$

Under the symmetric-response approximation, let $z_k^{\text{CTR}}$ denote the $k$th eigenvalue of $\mathbf{R}_{\text{CTR}}$, with corresponding normalized eigenvector $|\mathbf{x}_k\rangle$. The unperturbed eigenvalue of $c_R \mathbf{R}_{\text{CTR}}^{-1}$ is

$$\frac{c_R}{z_k^{\text{CTR}}}. \quad (78)$$

Using first-order perturbation theory, the corresponding eigenvalue of $\mathbf{R}_{\text{FR}}^{-1}$ is

$$\frac{1}{z_k^{\text{FR-like}}} \approx \frac{c_R}{z_k^{\text{CTR}}} + \mathbf{x}_k^T \mathbf{M} \mathbf{x}_k. \quad (79)$$

Because $\mathbf{D}_{\text{FR}}$ and $\mathbf{D}_{\text{CTR}}$ are diagonal, the perturbative correction can be written explicitly as

$$\mathbf{x}_k^T \mathbf{M} \mathbf{x}_k = \sum_{l=1}^N |x_{kl}|^2 \left[ \frac{1}{D_{l,\text{FR}}} - \frac{c_R}{D_{l,\text{CTR}}} \right]. \quad (80)$$

Thus,

$$\boxed{\frac{1}{z_k^{\text{FR-like}}} \approx \frac{c_R}{z_k^{\text{CTR}}} + \sum_{l=1}^N |x_{kl}|^2 \left[ \frac{1}{D_{l,\text{FR}}} - \frac{c_R}{D_{l,\text{CTR}}} \right]} \quad (81)$$

In practice, we approximate $\mathbf{D}_{\text{FR}} = \psi \mathbf{D}_{\text{CTR}}$ where $\psi$ is the ratio of the average susceptibility over all animals.

### 4.2 Consequences for the covariance spectrum and signal–noise alignment

For presynaptic fluctuations, the covariance eigenvalues are related to the response eigenvalues by

$$\lambda_k \sim (rg\sigma_\xi)^2 z_k^2. \quad (82)$$

Assuming that the amplitude of the presynaptic fluctuations is unchanged, the covariance-eigenvalue ratio is approximately

$$\frac{\lambda_k^{\text{FR}}}{\lambda_k^{\text{CTR}}} \simeq c_R^2 \left( \frac{z_k^{\text{FR}}}{z_k^{\text{CTR}}} \right)^2. \quad (83)$$

Thus, an increase in covariance eigenvalues despite the decrease in $rg$ implies that the corresponding response eigenvalues $z_k$ increase sufficiently to compensate for the reduced synaptic factor.

The changes in single-neuron gain are strongest for neurons on the shoulders of the stimulus-evoked responses, rather than for silent or highly active neurons. When propagated through the recurrent network, these heterogeneous changes increase the response eigenvalues on average, with smaller modes increasing more strongly than the largest modes. This explains the simultaneous increase in the average covariance eigenvalue and participation ratio: total variability increases, but the additional variance is spread across more modes rather than being concentrated only in the largest one.

We can also understand the changes in signal–noise alignment. Food restriction redistributes the signal across the altered response modes. The normalized overlap with covariance mode $k$ is

$$\cos^2 \phi_k = \frac{(\mathbf{x}_k^\top \Delta \boldsymbol{\mu})^2}{\|\Delta \boldsymbol{\mu}\|^2} = \frac{(z_k b_k)^2}{\sum_l (z_l b_l)^2}, \quad (84)$$

where

$$b_k = \mathbf{x}_k^\top \Delta \mathbf{I}. \quad (85)$$

If the eigenvectors and input projections $b_k$ are approximately preserved, the smaller relative increase of the largest noise modes reduces the overlap between the signal and these modes.

Overall, food restriction first alters single-cell gain. The recurrent network then propagates these heterogeneous gain changes through the population, shaping how signal and noise are distributed across response modes. The average covariance eigenvalue increases, intermediate modes are preferentially amplified, dimensionality increases, and the signal becomes less aligned with the largest noise modes. The result is a more variable but geometrically reorganised population code in which the loss of single-neuron selectivity can be partially compensated at the population level.

#### 4.3 Turning control tuning curves into food-restricted-like ones

To investigate the effects of food restriction on tuning-curve width, we inferred the effective input using the control data, transformed the linear-response eigenvalue distribution according to Eq. (81), and then reconstructed the tuning curves.

1. Calculate the covariance $\boldsymbol{\Sigma}^{\text{CTR}}$ of the control data.
2. Determine an effective linear response matrix $\mathbf{R}_0^{\text{CTR}}$ from the covariance:

$$\boldsymbol{\Sigma}^{\text{CTR}} = [(rg)_{\text{CTR}} \sigma_\xi]^2 \mathbf{R}^{\text{CTR}} (\mathbf{R}^{\text{CTR}})^\top = \mathbf{U} \text{diag} \left( \boldsymbol{\lambda}^{\text{CTR}} \right) \mathbf{U}^\top, \quad (86)$$

$$\mathbf{R}_0^{\text{CTR}} = \mathbf{U} \text{diag} \left( \frac{\sqrt{\boldsymbol{\lambda}^{\text{CTR}}}}{(rg)_{\text{CTR}} \sigma_\xi} \right) \mathbf{U}^\top, \quad (87)$$

where we choose the symmetric positive-semidefinite response matrix.

3. Using the measured control tuning curves, represented by the signal vector $\widehat{\Delta \boldsymbol{\mu}}^{\text{CTR}}$, infer the corresponding input vector:

$$\Delta \boldsymbol{\mu}^{\text{CTR}} = (rg)_{\text{CTR}} \mathbf{R}^{\text{CTR}} \delta \mathbf{I}, \quad (88)$$

$$\widehat{\delta \mathbf{I}}(\theta) = \frac{(\mathbf{R}_0^{\text{CTR}})^\top \widehat{\Delta \boldsymbol{\mu}}^{\text{CTR}}}{(rg)_{\text{CTR}}}. \quad (89)$$

4. Transform each control response eigenvalue using the above prescription Eq.81
5. We then apply the forward transformation to the signal vector:

$$\widehat{\Delta \boldsymbol{\mu}}^{\text{FR}} = (rg)_{\text{FR}} \mathbf{U} \text{diag} \left( \mathbf{z}^{\text{FR-like}} \right) \mathbf{U}^\top \widehat{\delta \mathbf{I}}. \quad (90)$$

We applied this procedure separately to all control animals and found that the transformed average tuning curve was broader. This shows that the flattening of the linear-response eigenvalue distribution induced by the inferred food-restriction parameters is sufficient to produce a widening of the tuning curves.

### 5 Mean Field Derivation

For each condition $q \in \{\text{CTR}, \text{FR}\}$, introduce an auxiliary binary activity variable $\sigma_i \in \{0, 1\}$. We approximate the joint distribution of the binary activities by the factorized variational distribution

$$Q_{\rho}(\boldsymbol{\sigma}) = \prod_{i=1}^N \rho_i^{\sigma_i} (1 - \rho_i)^{1-\sigma_i}, \quad (91)$$

where

$$\rho_i = \langle \sigma_i \rangle_{Q_{\rho}} \quad (92)$$

is the mean activity of neuron $i$.

The entropy of this factorized distribution is

$$S[Q_{\rho}] = - \sum_{i=1}^N [\rho_i \log \rho_i + (1 - \rho_i) \log(1 - \rho_i)]. \quad (93)$$

Using the mean-field replacement $\sigma_i \sigma_j \mapsto \rho_i \rho_j$, the variational free energy is

$$\begin{aligned} \mathcal{F}_q(\boldsymbol{\rho}; \theta) = & -\frac{c_q}{2} \boldsymbol{\rho}^T \mathbf{W} \boldsymbol{\rho} + \frac{c_q w_0}{2N} \left( \sum_i \rho_i \right)^2 - c_q \sum_i (I_i(\theta) - \bar{I}(\theta)) \rho_i - b_q \sum_{i=1}^N \rho_i \\ & + T \sum_{i=1}^N [\rho_i \log \rho_i + (1 - \rho_i) \log(1 - \rho_i)], \end{aligned} \quad (94)$$

where $\bar{I}$ is the average input over the neurons and the translational-invariant CANN connectivity has the same sum for every column and row $\sum_j W_{ij} = w_0$. The derivative of the entropy contribution is

$$\frac{\partial}{\partial \rho_i} [\rho_i \log \rho_i + (1 - \rho_i) \log(1 - \rho_i)] = \log \left( \frac{\rho_i}{1 - \rho_i} \right). \quad (95)$$

Consequently,

$$\frac{\partial \mathcal{F}_q}{\partial \rho_i} = -c_q \left[ (\mathbf{W} \boldsymbol{\rho})_i + I_i(\theta) - \bar{H} \right] - b_q + T \log \left( \frac{\rho_i}{1 - \rho_i} \right). \quad (96)$$

where we define the population average

$$\bar{H} = \frac{1}{N} \sum_{j=1}^N [(\mathbf{W} \boldsymbol{\rho})_j + I_j(\theta)]. \quad (97)$$

Setting Eq. (96) to zero gives

$$T \log \left( \frac{\rho_i}{1 - \rho_i} \right) = c_q \left[ (\mathbf{W} \boldsymbol{\rho})_i + I_i(\theta) - \bar{H} \right] + b_q. \quad (98)$$

Solving for $\rho_i$ yields

$$\rho_i = \frac{1}{1 + \exp \left\{ -\frac{c_q}{T} \left[ (\mathbf{W}\boldsymbol{\rho})_i + I_i(\theta) - \overline{H} \right] - \frac{b_q}{T} \right\}}. \quad (99)$$

Finally, define the logistic transfer function

$$F(v) = \frac{1}{1 + \exp(-v)} \quad (100)$$

and identify the dimensionless parameters used in the Methods as

$$rg = \frac{c_q}{T}, \quad b = \frac{b_q}{T}. \quad (101)$$

The mean-field self-consistency equation is then

$$\boxed{\rho_i = F \left[ rg \left( (\mathbf{W}\boldsymbol{\rho})_i + I_i(\theta) - \overline{H} \right) + b \right]}, \quad (102)$$

which is Eq. (14) in the Methods.

### 6 Temporal Inference

We infer the temporal change in input gain by comparing how the signal projection and covariance eigenvalue change along each response mode. We introduce a gain $\gamma$ that modulates the stimulus-dependent mean input. This modelling choice is motivated by evidence that mean responses and trial-to-trial variability can be modulated separately: stimulus onset can reduce cortical response variability even when mean firing rates change little<sup>21</sup>. The covariance eigenvalue, defined above is given as

$$\lambda_k = \sigma_\xi^2 c^2 z_k^2 + \frac{z_k}{L}. \quad (103)$$

Under the symmetric-response approximation, let

$$\mathbf{R}\mathbf{x}_k = z_k \mathbf{x}_k, \quad (104)$$

where $z_k$ is the response eigenvalue associated with mode $\mathbf{x}_k$. The projection of the signal vector onto this mode is

$$s_k = \mathbf{x}_k^\top \Delta \boldsymbol{\mu}. \quad (105)$$

Using

$$\Delta \boldsymbol{\mu} = c\gamma \mathbf{R}\Delta \mathbf{I}, \quad c = rg, \quad (106)$$

this projection can be written as

$$s_k = c\gamma z_k b_k, \quad b_k = \mathbf{x}_k^\top \Delta \mathbf{I}. \quad (107)$$

We define the ratios between the food-restricted and control conditions as

$$a_k = \frac{s_k^{\text{FR}}}{s_k^{\text{CTR}}}, \quad z_{k,R} = \frac{z_k^{\text{FR}}}{z_k^{\text{CTR}}}, \quad \lambda_{k,R} = \frac{\lambda_k^{\text{FR}}}{\lambda_k^{\text{CTR}}}, \quad (108)$$

together with

$$\gamma(t)_R = \frac{\gamma^{\text{FR}}(t)}{\gamma^{\text{CTR}}(t)}, \quad c_R = \frac{c^{\text{FR}}}{c^{\text{CTR}}} = \frac{(rg)^{\text{FR}}}{(rg)^{\text{CTR}}}. \quad (109)$$

We assume that the stimulus input $\Delta \mathbf{I}$ is unchanged between conditions and that corresponding response modes can be matched between the control and food-restricted conditions. The input projection $b_k$ then cancels when taking the ratio of Eq. (107), giving

$$a_k = c_R \gamma(t)_R z_{k,R}. \quad (110)$$

The change in the response eigenvalue can therefore be written as

$$z_{k,R} = \frac{a_k}{c_R \gamma(t)_R}. \quad (111)$$

To express the covariance ratio, we define

$$P_k = L \sigma_\xi^2 c_{\text{CTR}}^2 z_k^{\text{CTR}}. \quad (112)$$

This dimensionless parameter is the ratio between the slow and fast contributions to the control covariance along mode $k$. Dividing the FR and control versions of Eq. (103) gives

$$\lambda_{k,R} = \frac{P_k c_R^2 z_{k,R}^2 + z_{k,R}}{P_k + 1}. \quad (113)$$

Substituting Eq. (111) then gives

$$\lambda_{k,R} = \frac{P_k \frac{a_k^2}{\gamma(t)_R^2} + \frac{a_k}{c_R \gamma(t)_R}}{P_k + 1}. \quad (114)$$

Equation (114) relates the measured change in covariance, $\lambda_{k,R}$, to the measured change in the signal projection, $a_k$, and the unknown gain ratio $\gamma(t)_R$. The two covariance contributions depend differently on the gain.

We estimate $\gamma(t)_R$ by fitting Eq. (114) jointly across response modes. The parameters $P_k$ are treated as mode-dependent but constant over time, while $\gamma(t)_R$ is shared across modes within each time bin. The coupling ratio $c_R$ is fixed to the value inferred in Extended Data Fig. 8a. This separates the time-dependent change in external gain from the time-independent changes in cellular and recurrent response.
